# Genetically engineered microglia-like cells have therapeutic potential for neurodegenerative disease

**DOI:** 10.1101/2021.09.16.460703

**Authors:** Robert N. Plasschaert, Mark P. DeAndrade, Fritz Hull, Quoc Nguyen, Tara Peterson, Aimin Yan, Mariana Loperfido, Cristina Baricordi, Luigi Barbarossa, John K. Yoon, Yildirim Dogan, Zeenath Unnisa, Jeffrey W. Schindler, Niek P. van Til, Luca Biasco, Chris Mason

**Affiliations:** AVROBIO, Inc., Cambridge, MA, USA; Child Neurology, Emma Children’s Hospital, Amsterdam University Medical Centers, Vrije Universiteit and Amsterdam Neuroscience, Amsterdam, the Netherlands; Great Ormond Street Institute of Child Health, University College London; Advanced Center for Biochemical Engineering, University College London

## Abstract

Hematopoietic stem/progenitor cell gene therapy (HSPC-GT) results in the engraftment of genetically modified microglia-like cells (MLCs) in the brain. While HSPC-GT has shown a clear neurological benefit in the clinic for specific rare diseases, the nature of MLC engraftment in the brain and the functional characteristics of MLCs remain contentious. Here we comprehensively characterized how different routes of administration affect the engraftment and biodistribution of genetically engineered HSPC-derivatives in mice. Using high-throughput single-cell profiling, we show that MLCs bear a transcriptional signature similar to resident microglia rather than invading macrophages. However, MLCs could clearly be distinguished from resident microglia by expression of a specific set of genes. Finally, in murine models of Parkinson’s disease and frontotemporal dementia, we demonstrate that MLCs can provide therapeutically relevant levels of protein to the brain, thereby potentially opening avenues of HSPC-GT to address the underlying disease etiology of these and other similar disorders.

## Introduction

Hematopoietic stem/progenitor cell gene therapy (HSPC-GT) is a well-established paradigm for treating monogenic diseases, with more than 300 patients currently dosed^1-4^. Clinical applications of HSPC-GT include treatments of inherited immune deficiencies (e.g., SCID-X1, ADA-SCID, Wiskott-Aldrich syndrome), blood disorders (e.g., sickle cell disease, transfusion-dependent beta-thalassemia, X-linked chronic granulomatous disease), and lysosomal storage disorders (e.g., Hurler syndrome, Hunter syndrome, Fabry disease)^5-13^. In general, a patient’s bone marrow cells are mobilized and collected from the periphery and HSPCs are isolated and transduced ex vivo with a lentiviral vector carrying a therapeutic payload. The genetically engineered cells are then intravenously infused back into the patient after a conditioning regimen using a myeloablative agent, such as busulfan. In mice, non-human primates, and humans, gene-modified HSPCs can engraft long-term throughout the body, including within the hematopoietic compartment as cells of the bone marrow and blood, within the peripheral organs as tissue-resident macrophages, and within the brain and spinal cord as microglia-like cells (MLCs)^14-17^.

The migration of genetically engineered cells across the blood-brain barrier (BBB) and their engraftment within the brain parenchyma is key to HSPC-GT’s ability to address neurological dysfunction. The conditioning agent regime causes a partial depletion of microglia that can be repopulated by genetically modified MLCs that produce a therapeutic protein^18^. Additionally, these MLCs can secrete this therapeutic protein and thereby provide a local source for uptake by neighboring cells, such as neurons, astrocytes, and oligodendrocytes. Late-stage HSPC-GT clinical trials for metachromatic leukodystrophy (MLD) and cerebral adrenoleukodystrophy (CALD) have demonstrated halting of expected CNS-related disease progression, with most patients being free of severe motor and cognitive dysfunction^19-21^. The fact that HSPC-GT can address the neurological symptoms of rare genetic diseases supports its possible translation to more common neurodegenerative disorders, such as Parkinson’s disease and dementia.

Still, a wider application of HSPC-GT to neurological diseases requires a deeper understanding of the nature and dynamics of genetically engineered MLCs and their engraftment in the brain. This includes the extent and durability of their engraftment within the CNS, the effect of different routes of HSPC administration, and the comparability of gene-modified MLCs to endogenous microglia, given that these two populations are developmentally distinct. In the present study we have addressed these key questions in preclinical murine models of HSPC-GT and demonstrate that this platform can provide expression of potentially therapeutic transgene at a potentially therapeutic level throughout the periphery and the brain in two mouse models of neurodegenerative disease.

## Results

### Intravenous (IV) and intracerebroventricular (ICV) administration both show engraftment in the brain but only IV shows engraftment in the periphery

Work in murine disease models have shown evidence of engraftment and functional rescue in the periphery and the brain after HSPC-GT using intravenous (IV) administration^22-28^. It was also previously shown that intracerebroventricular (ICV) administration of genetically engineered HSPCs directly into the brain results in widespread engraftment of cells with many characteristics of endogenous microglia^18^. However, the extent of engraftment and durability of HSPC-derived cells in the brain and how the route of administration effects the nature of engrafted cells are poorly understood. To shine light on these crucial aspects, we performed a comprehensive animal biodistribution study following IV, ICV, or a combination of IV and ICV dosing of genetically engineered HSPCs into C57BL/6J mice after busulfan conditioning (Fig. 1a).

**Figure 1.**
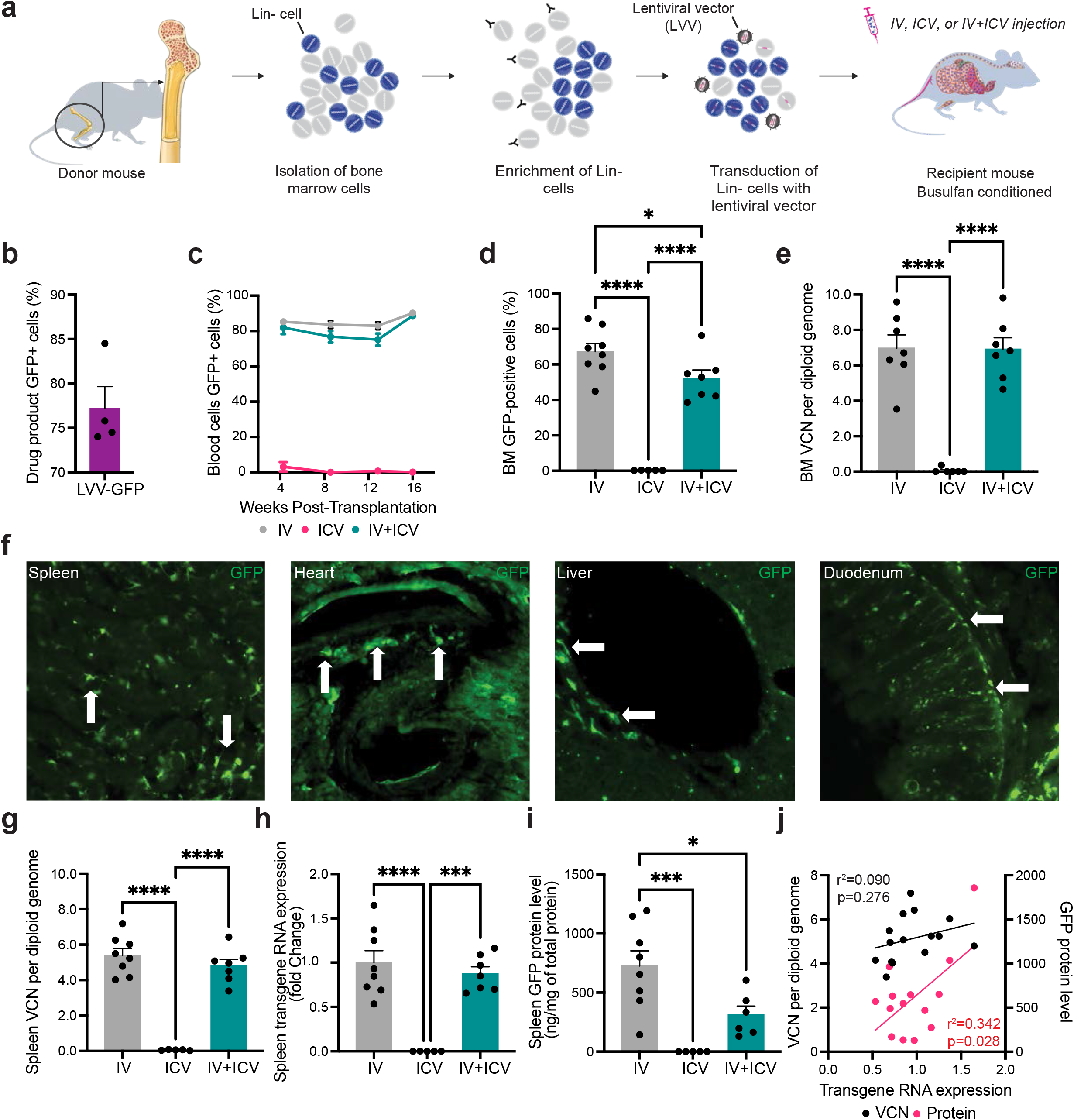
Peripheral engraftment HPSC-derived cells following administration via IV, ICV or IV+ICV. **a** Total bone marrow cells were isolated from the long bones of donor mice, enriched for lineage negative (Lin-) hematopoietic stem/progenitor cells and transduced with a lentiviral vector that drives the expression of GFP (LVV.GFP). Busulfan conditioned recipient animals received cell doses via IV, ICV, or both (IV+ICV). **b-d** Flow cytometry for GFP was performed on the transduced Lin-drug product, at four-week intervals on peripheral blood, and on the bone marrow (BM) at necropsy (16 weeks post-transplantation). **e** Vector copy number (VCN) in the bone marrow was determined by qPCR. **f** Peripheral organs were harvested, fixed, sectioned, and GFP-positive cells were observed in all organs examined for all IV-dosed animals examined, including the spleen, heart, liver, and duodenum. No such engraftment was observed in any dosed via ICV-alone. **g-i** VCN, transgene RNA expression, and GFP protein levels were measured in spleens from each treatment group at 16 weeks post treatment. **j** Correlation analysis between splenic levels of RNA and VCN (black dots and bar) and splenic levels of RNA and protein (red dots and bar) for animals that received doses by IV or IV+ICV. Bars represent means ± Standard Error of the Mean (SEM). Closed circles in each graph represent individual data points. For c-e and g-I, a one-way ANOVA with Tukey’s multiple post-hoc comparison test was conducted. For j, the Pearson correlation coefficient for each correlation is shown.

Lineage negative (Lin-) HSPCs were isolated (Supp. Fig. 1a, b) and transduced with a lentiviral vector encoding GFP (LVV.GFP) at greater than 74% transduction efficiency (range: 74.0%-84.5% GFP-positive cells; Fig. 1b) and approximately 6 copies of integrated vector per genome (Supp Fig. 1c). No changes to the proliferative potential of the drug product were observed after transduction, as measured using the colony forming unit assay (Supp. Fig. 1d). Recipient animals received 4 days of 25 mg/kg/q.d. (cumulative dose of 100 mg/kg) of busulfan prior to cell administration and lost 10-15% of their bodyweight which was rapidly regained post-treatment (Supp. Fig. 1e). As expected, robust chimerism of all major cell types in the peripheral blood and bone marrow was observed in animals that received an IV dose of genetically engineered HSPCs, but not in animals that only received genetically engineered HSPCs via ICV (Fig. 1c-e, Supp. Fig. 2). Similarly, engraftment of cells throughout the peripheral organs was specifically limited to animals that received an IV dose (Fig. 1f). In the spleen, we detected integrated vector (Fig. 1g), transgene expression (Fig. 1h), and GFP protein expression (Fig. 1i) at 16 weeks after infusion only in animals that received cells via an IV dose. As expected, we observed a significant correlation between the levels of GFP transcript and total amount of GFP protein in the periphery (Fig. 1j).

Within the brain, all three routes of administration led to widespread engraftment of GFP-positive cells throughout the rostral-caudal axis, including the cortex, hippocampus, choroid plexus, thalamus, and cerebellum. We assessed engraftment at 16 weeks post-transplantation (Fig. 2a, b) and then again at 12-13 months, observing widespread engraftment at both timepoints. At 16 weeks, 78-89% of GFP+ cells from all routes of administration co-expressed the microglial marker Iba1 (IV: 78.40% ± 3.30%, ICV: 88.67% ± 2.78%; IV+ICV: 81.57% ± 3.20%; Fig. 2c). Moreover, GFP-positive cells that colocalized with Iba1 had a ramified morphology consistent with the physiological surveilling behavior of endogenous microglia (Fig. 2a). Strikingly and in contrast to previous reports^18^, we observed a significant difference in the level of engraftment between the IV and ICV routes of administration. Animals dosed by IV alone had an average of 19.02% (range: 13.04%-25.44%) microglia replaced compared to an average of 5.71% (range: 1.87%-8.26%) microglia replaced in animals dosed by ICV alone (Fig. 2d). Animals dosed using both routes of administration (IV+ICV) had a similar level to the IV alone dosed animals (range: 14.98%-26.64%; mean: 19.35% ± 1.57%; Fig. 2d). Measurements of vector copy number (VCN) per genome (Fig. 2e), transgene expression (Fig. 2f), and GFP protein levels in the whole brain (Fig. 2g) corroborated this difference in engraftment between IV and ICV dosing. A statistically significant correlation was found between VCN and GFP mRNA and GFP protein levels and GFP mRNA in the brain (Fig. 2h). Of note, we observed a general lack of correlation of biodistribution metrics (VCN, mRNA, protein) in peripheral tissues versus brain, consistent with previous clinical and preclinical reports^15,20,29^ that engraftment in these two compartments are independent events that occur from separate cell populations derived from the drug product (Supp. Fig. 3).

**Figure 2.**
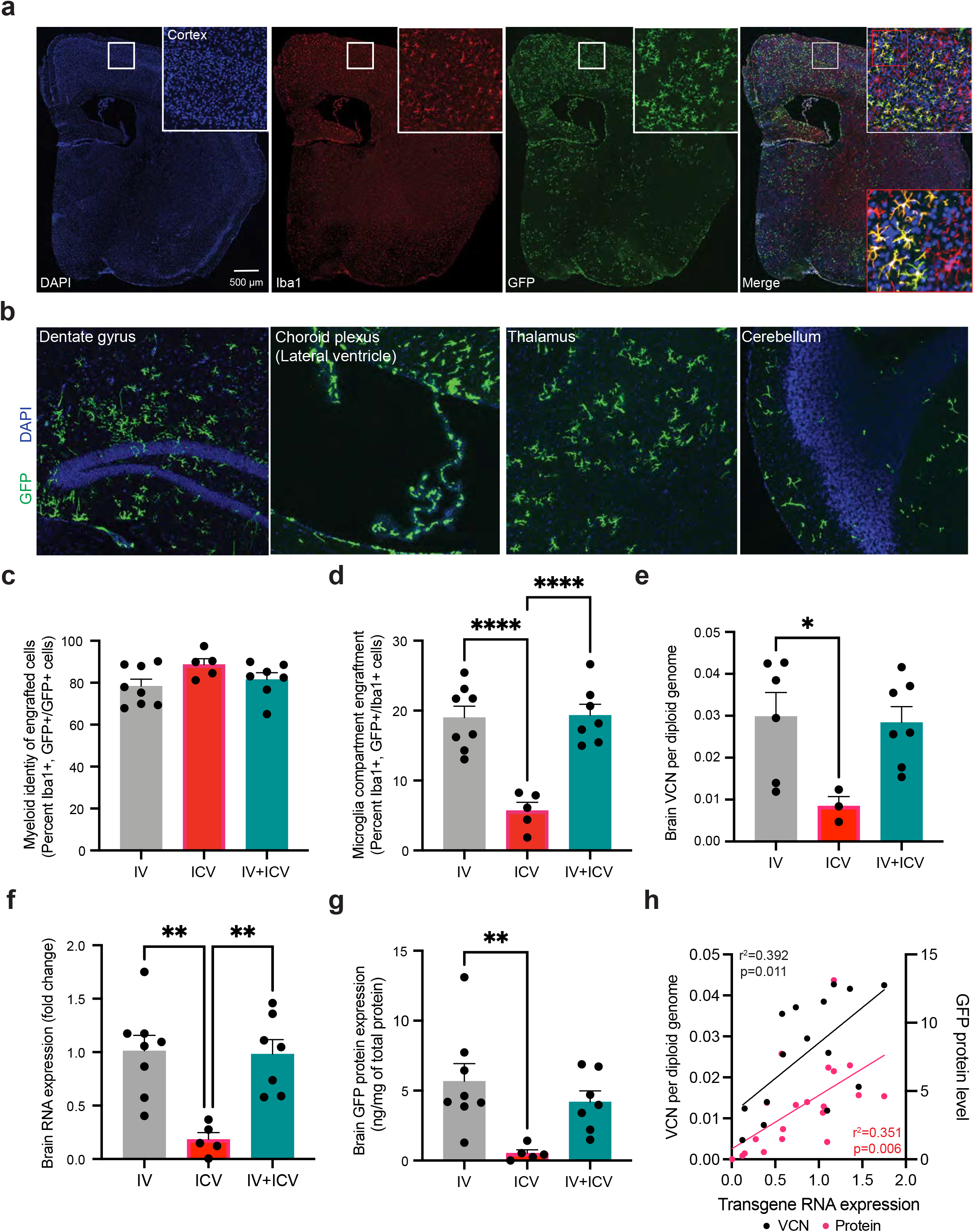
Engraftment of MLCs in the brain following different routes of administration of genetically engineered HSPCs. **a** Representative image of a coronal slice from an IV-dosed animal visualizing Iba1, DAPI and GFP at 16 weeks post-transplantation. Top right insert surrounded by white box is a digital magnification of a region of the primary motor cortex from the whole brain. Bottom right insert surrounded by red box is a magnification of the image in the white box. **b** Representative images from an IV-dosed animal at 16-weeks post-transplantation with GFP-positive cells located throughout the rostral-caudal axis of the brain, including the hippocampus, choroid plexus, thalamus, and cerebellum. **c** Quantification of GFP-positive engrafted cells that express the myeloid marker Iba1. **d** Quantification of Iba1+ cells that expressed GFP. **e-g** Quantification of integrated vector into the genome (VCN) via qPCR, RNA transgene expression via RT-qPCR, and GFP protein levels via ELISA in the brain for all treatment groups. **h** Correlation analysis of transgene RNA expression and VCN and transgene RNA expression and protein levels. Error bars represent means ± SEM. For c-g Tukey’s multiple post-hoc comparison test was conducted. For h, the Pearson correlation coefficient for each correlation is shown.

### HSPC-derived MLCs are highly similar but are transcriptionally distinguishable from endogenous microglia

The characteristics of HSPC-derived MLCs that engraft in the brain and their equivalence to microglia, which are derived from the embryonic yolk sac, remain contentious. Differences in the source of bone marrow-derived progenitors (e.g., the bone marrow compartment versus an administered cell bolus), the nature of cell recruitment (e.g., native signaling versus induction from microglial ablation), and the state of the brain (e.g., healthy versus active disease), all influence MLCs and how they might compare to *bona fide* microglia. While transcriptional signatures that separate HSPC-derived MLCs and microglia have been described broadly, the single cell heterogeneity of these populations and a high-resolution characterization of these differences in HSPC-GT have remained uncovered to date. Similarly, the transcriptional signatures of long-term engrafted MLCs that engraft via migration from the periphery (IV-dosed) versus those directly administered to the brain (ICV-dosed) have not been characterized.

To address these aspects, we combined cell sorting with high-throughput single-cell RNA (scRNA) analyses to characterize long-term engrafted MLCs and endogenous microglia from the same animals. We isolated endogenous microglia (GFP-negative) and MLCs (GFP-positive) from enzymatically dissociated whole brains by FACS based on the expression of cell surface markers (CD45+, CD11b+, CX3CR1+) from one male and one female mouse 12-13 months after either IV or ICV administration of HSPCs. After sequence processing and data analysis, we obtained a total of 29,085 individual scRNA profiles across the eight combined datasets for MLCs and endogenous microglia. We also generated comparator datasets from a 12-week-old, treatment naïve C57BL/6J animal for several cell populations: microglia (CD45+, CD11b+, CX3XCR1+), neurons (CD45-, Thy1+, ACSA2-), astrocytes (CD45-, Thy1-, ACSA2+) and peripheral blood mononuclear cells (PBMCs; Fig 3a.; Supp. Fig. 4). By tSNE dimensionality reduction of the most variable genes, we observed that MLCs are in distinct but neighboring clusters to endogenous microglia (Fig 3b, c; Supp. Fig. 5). The majority of PBMCs, neurons, and astrocytes occupied largely distinct single cell clusters as compared to microglia and MLCs, except notably a population of PBMC circulating monocytes which cluster with MLCs. Hierarchical clustering analysis confirmed that MLCs are similar to endogenous microglia and share higher similarity with bone marrow-derived PBMCs than neurons and astrocytes (Fig. 3d, e). We then directly compared the normalized levels of individual transcripts in all GFP-positive MLCs to that of their endogenous GFP-negative microglia counterparts to assess the degree of similarity between these two populations. We observed a strong correlation across most transcripts (Fig. 3f; adjusted R^2^= 0.9071), supporting that the transcriptome of HSPC-derived MLCs are distinguishable, but largely similar to that of endogenous microglia.

**Figure 3.**
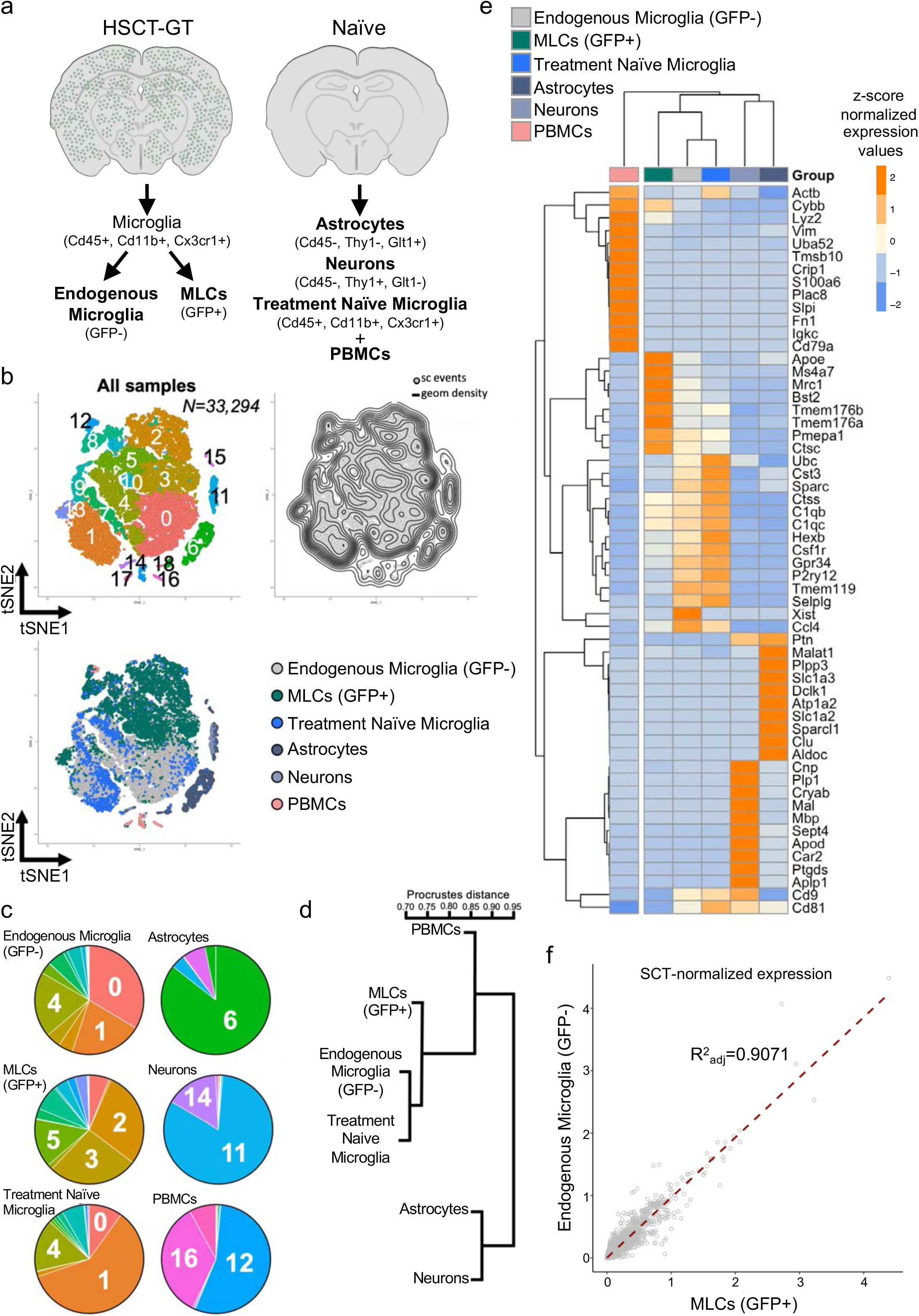
Single-cell comparison of MLCs, endogenous microglia, PBMCs, neurons, and astrocytes. **a** MLCs and endogenous microglia were isolated via enzymatic digestion, FACS-purified, and then single cell sequenced from IV and ICV dosed male and female animals. PBMCs, neurons, astrocytes, and microglia were similarly isolated from a treatment naïve animal. **b** Global tSNE plot generated from a total of 33,294 cells (n=5 mice; 12 samples). The plot on the left shows the Louvain clustering (resolution 0.4) while the plot on the right shows the density of single cell events on the map. **c** Pie charts showing the cluster breakdown by sample, with labels showing the top represented clusters for each sample type. 7 MLC-enriched clusters (2-5,8,10,12,13,15), 5 microglia-enriched clusters (0,1,4,7,9), 1 neuron cluster (11), 1 astrocyte cluster (6) and 4 PBMC clusters (14, 16-18) were identified. **d** Unsupervised clustering based on the global expression profile of each sample type. **e** Heatmap showing top 10 differentially expressed genes for each sample type. Dendrogram at the top shows samples clustering based on expression profiles of these genes. **f** Correlation analysis of endogenous microglia versus MLCs based on normalized single cell gene expression data.

We then analyzed in detail the single cell heterogeneity of GFP-positive MLCs and GFP-negative endogenous microglia in our treated mice. By tSNE dimensionality reduction of highly variable genes and Louvain clustering of single cells, we observed 8 main single-cell clusters when comparing these two populations. Notably, we observed a clear separation of MLC and microglia populations using clustering analysis (Fig. 4a, b; Supp. Fig. 6). GFP-negative endogenous microglia were enriched in clusters 1,2,4, and 6, while GFP-positive MLCs are enriched in clusters 0, 3, and 5. Clusters 7 and 8 are composed of a very small number of contaminating glia (astrocytes and oligodendrocytes) which we preserved as an outgroup comparison. The top 10 significantly enriched genes defining each cluster highlight the developmental differences between MLCs and microglia and potential differences in inflammatory state (Fig. 4c, d). Clusters containing mostly GFP-negative cells were enriched for markers associated with microglia homeostasis including *Mef2a, Hexb, Ctsc3, P2ry12, Selplg*, and *Sparc*, with this signature being most highly enriched in clusters 2 and 6. Clusters 1 and 4 show relatively lower levels of homeostatic genes, a hallmark of mild microglia activation that is likely in part a result of the enzymatic isolation process. Notably, cluster 1 clearly separates from cluster 4 based almost entirely on the sex of the recipient animal, with differences driven largely by the expression of the X-inactivation genes *Xist* and *Tsix*. This sex difference was not observed in GFP-positive MLC-enriched clusters, given that all donor animals for this experiment were males (Supp. Fig. 7). Clusters containing mostly GFP-positive MLCs were enriched with core transcriptional markers previously associated with bone-marrow derived, ‘CNS associated macrophages’ (CAMs) *Ms4a7, Mrc1, Pf4*, and *Stab1*^*30*^. Moreover, GFP-positive enriched clusters also show stronger expression of activation markers of endogenous microglia including genes associated with reactive oxidation (*Cybb, Fyb;* cluster 0), metabolism (*Apoe, Lyz2*; cluster 3), and genes that encode for the major histocompatibility complexes (MHC) such *Cd74, H2-Aa, H2-Eb1*, and *H2Ab1* (cluster 5).

**Figure 4.**
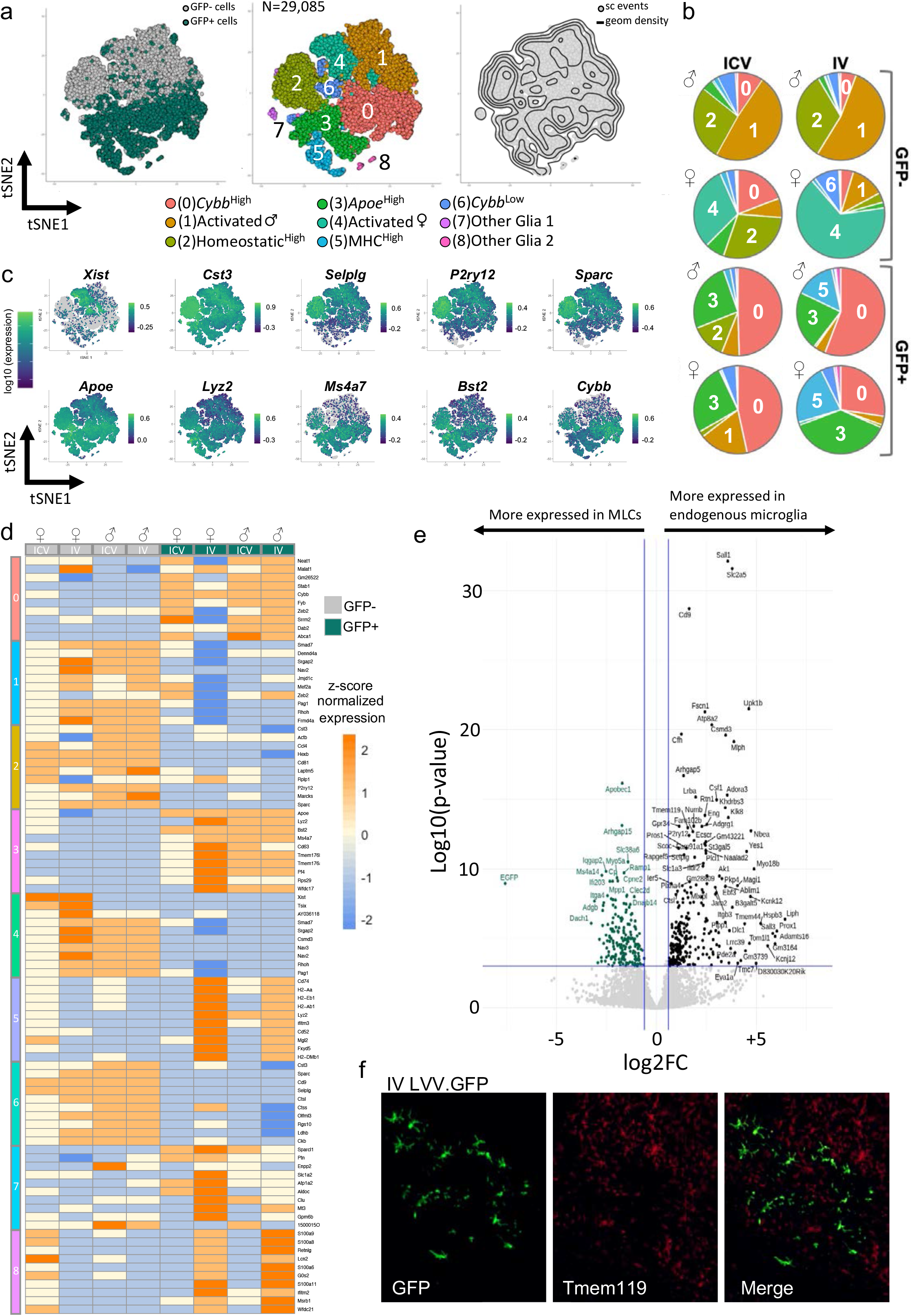
Characterization of the single-cell transcriptional landscape of MLCs versus endogenous microglia in treated mice. **a** Global tSNE plot generated from a total of 29,085 single cells from endogenous microglia (GFP) and MLCs (GFP+, in green) (n=4 mice; 8 samples). The plot on the left shows single cells separated in the main GFP-negative (gray dots) and GFP-positive (green dots) groups. The plot in the middle shows the Louvain clustering (resolution 0.2). Putative functional associations and/or key markers differentiating each cluster are listed below. The plot on the right shows the density of single cell events in the map. **b** Feature plots showing expression of individual genes characterizing the top (microglia enriched) or the bottom (MLCs enriched) sections of the map. **c** Heatmap showing single-cell normalized expression of the top 10 differentially expressed for each cluster as compared to the rest of the map (note: clusters might share top represented genes). Columns represent GFP-negative (grey squares) and GFP-positive (green squares) cells from treated mice labeled according to their sex, identification number and route of administration (ICV, intracerebroventricular; IV, intravenous). **d** Volcano plot highlighting differentially expressed genes between GFP-positive MLCs (left hand side highlighted in green) and GFP-negative microglia (x-axis log2 fold change in average gene expression, y-axis log. **e** Representative IHC images of GFP and Tmem119 from an IV-dosed animal.

To identify highly expressed markers that could distinguish MLCs from endogenous microglia, we first performed a differential gene expression analysis comparing the composite single-cell transcriptome of GFP-positive and GFP-negative cells (Fig. 4e). Homeostatic genes used as canonical microglia markers including *SallI, Csf1*, and *Tmem119* were enriched in GFP-negative endogenous microglia over GFP-positive MLCs. Accordingly, Tmem119 expression was evident in endogenous microglia but largely excluded from GFP-positive cells in brain (Fig. 4f). Genes enriched in MLCs include *Apobec1* and *Ms4a14*, markers that have been previously associated with BM-derived, CNS/PNS-associated macrophages when compared to endogenous microglia^31,32^. Gene Ontology (GO) and pathway analysis comparing the two populations highlight differences in genes associated with immune function, including chemokine production, cell adhesion, and responses to interferon, which is consistent with the functional role of microglia and MLCs in the brain (Supp. Figure 8).

Our analysis comparing MLCs and microglia confirms and expands the information from previous studies on CNS/PNS-associated macrophages, where expression of markers associated with inflammation in parenchymal microglia are enriched in bone marrow-derived macrophages isolated from healthy animals without overt neuroinflammation^29,32^. In addition, our data show clear concordance with markers previously identified using bulk RNA-Seq comparing MLCs and microglia after busulfan conditioning and IV dosing of HSPCs (Fig. 5a)^29^. We then compared our datasets to previous work characterizing the transcriptional profiles of CNS-invading, inflammatory macrophages that are mobilized from the periphery in the context of disease or injury^33^. Our comparison using this gene-list associated with invading macrophages showed that our GFP-positive MLCs show generally low expression of genes associated with invading macrophages and relatively higher expression of the homeostatic microglia signature (Fig. 5b). These results suggest that certain differences we observe between MLCs and microglia may result from their distinct developmental origin rather than from distinct activation states.

**Figure 5.**
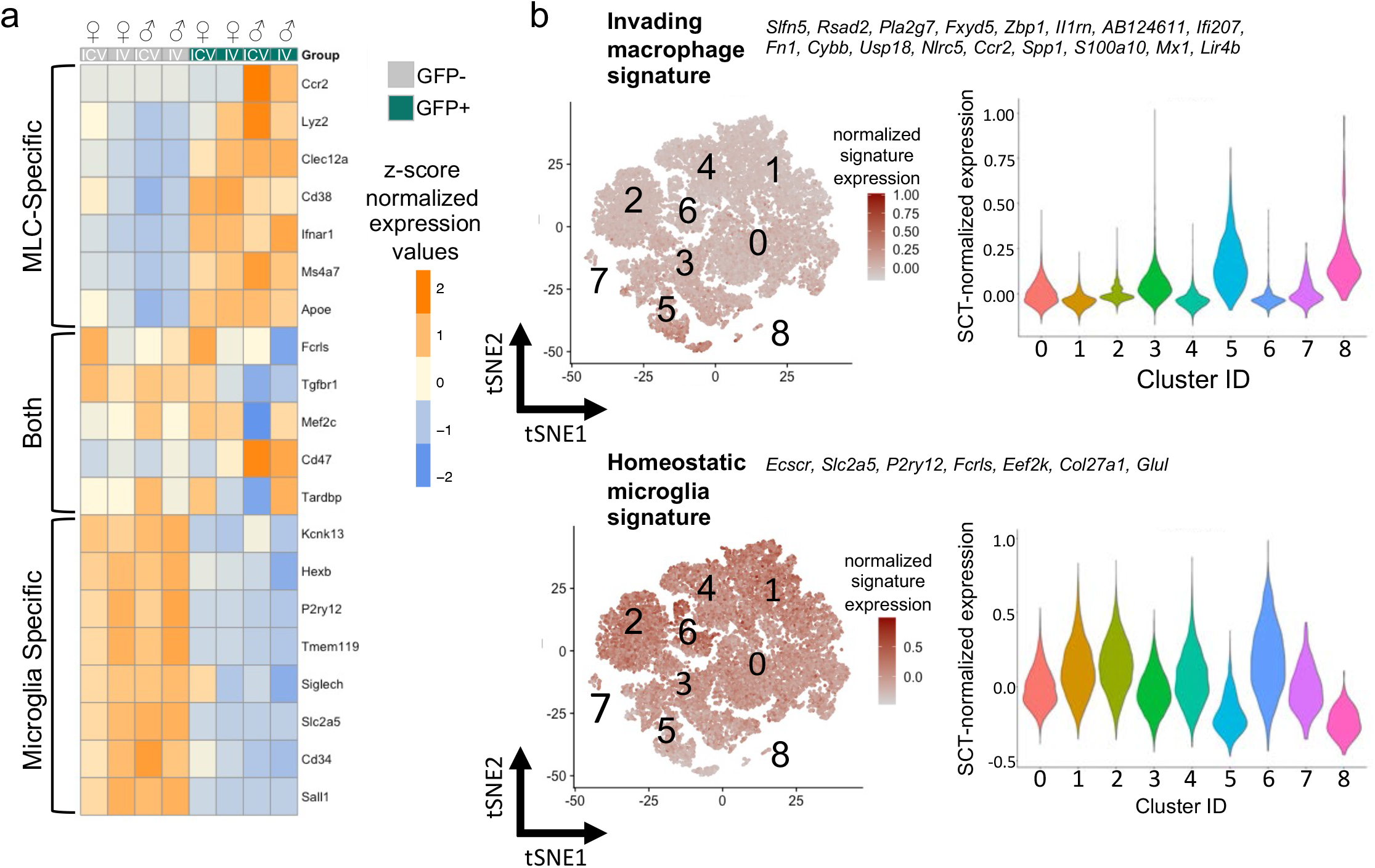
Projection of different MLC- and microglia-specific markers on the single-cell landscape. **a** Heatmap showing the single-cell normalized average expression values of MLC-specific, microglia specific and shared gene markers as described in Shemer *et al*., 2018 in GFP-negative and GFP-positive cells from treated mice. **b** Projection of invading versus homeostatic microglia gene signatures as described in DePaula-Silva *et al*., 2019 over the tSNE map of Fig 4a (left panels) and violin plots showing the expression of these signatures in the clusters identified in Fig.4a (right panels). **c** Violin plots showing the expression of genes associated with invading macrophages/monocytes (left panels) or homeostatic microglia (right panels) as described in Haage *et al*., 2019 in GFP-negative (grey squares) or GFP-positive (green squares) treated mice labeled according to their sex and identification number.

Lastly, we wanted to investigate whether the route of administration impacted the transcriptional profile of MLCs. Surprisingly and despite a vastly different engraftment history, we observed a significant overlap in the transcriptional signatures of IV-dosed GFP-positive and ICV-dosed GFP-positive cells, with most transcripts showing strong correlation in expression (Supp. Fig. 9a; R^2^=0.9134). IV and ICV GFP-positive cells had similar representation in each cluster, apart from cluster 5 (MHC^high^), which was enriched for IV GFP-positive cells over ICV GFP-positive cells (Fig. 4b). Expression of specific MHC genes was previously reported to be associated with so-called border-associated macrophages (BAMs), which are bone marrow-derived macrophages that occupy a niche near the blood-brain barrier (BBB)^34^. This signal is consistent with the engraftment history of IV-derived MLCs, which must migrate across the BBB. Accordingly, clustering analysis show differences between IV and ICV groups in the expression of several MHC genes, including *H2-a, H2-b*, and *Cd74* (Supp. Fig 9b)^35^.

### LVV.GBA and LVV.GRN increase GCase/progranulin expression and secretion

Our characterization of MLCs and our assessment of biodistribution of gene-modified cells supports a wider application of HSPC-GT to address the peripheral and CNS components of neurodegenerative disease with well-defined genetic risk factors. Two such disorders are GBA-associated Parkinson’s disease (GBA-PD) and progranulin-associated frontotemporal dementia (GRN-FTD), both of which are associated with the heterozygous loss of function of a lysosomal protein (beta glucocerebrosidase and progranulin, respectively). We generated lentiviral vectors expressing either codon optimized human GBA (LVV.GBA) or codon optimized human GRN (LVV.GRN) (Fig 6a) and transduced mouse macrophage RAW264.7 cells with increasing amounts of vector. Increased integrated vector copies per diploid genome predictably increased the amount of transgene-derived RNA transcript and protein expression for both vectors (Fig. 6b-q). Importantly, we observed active beta-glucocerebrosidase (GCase) and human progranulin (hGRN) in the conditioned media even at low VCN, supporting that lentiviral-mediated, supraphysiological expression of these proteins results in their secretion from macrophage-like cells (Fig. 6e, g, m, o). This suggests that GCase or progranulin could be secreted by microglia-like cells for uptake by other cells within the brain.

**Figure 6.**
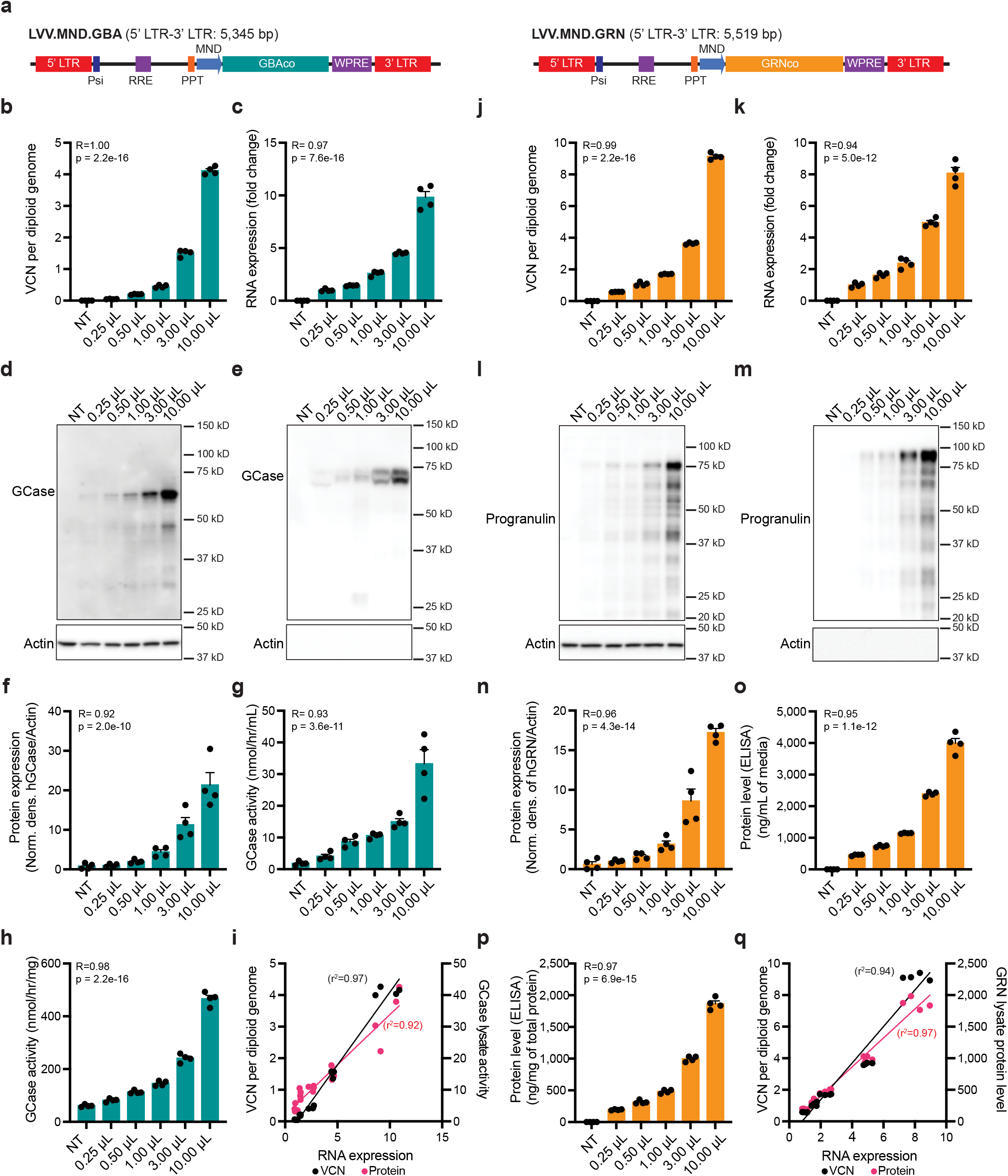
Characterization of lentiviral vectors for GBA and GRN in a mouse macrophage cell line. **a** Schematic of integrating component of lentiviral vectors from the 5’ long terminal repeat (LTR) to 3’ LTR. **b-i** LVV-GBA increases integrated vector copy number (VCN), transgene RNA expression, GCase protein expression, and GCase enzymatic activity levels in the lysate and conditioned media. **j-p** LVV-GRN increases integrated VCN, transgene RNA expression, and progranulin protein levels in the lysate and conditioned media. Bars represent means ± SEM. Pearson correlation analysis is shown where indicated.

### HSPC-GT in a model of GBA-PD results in widespread distribution of gene-modified cells in key tissues

Mutations in *GBA* are causative for the lysosomal storage disorder Gaucher disease and are among the most prevalent genetic risk factors for Parkinson’s disease^36,37^ and Lewy body dementia^38^. To assess the utility of lentiviral HSPC-GT for Parkinson’s disease patients associated with a *GBA* mutation (GBA-PD), we conducted a biodistribution study of key organs in homozygous *Gba*^D409V^ mice, a model of GBA-PD^39^. We isolated lineage negative cells and transduced them with either a lentiviral vector encoding our previously characterized human GBA transgene (LVV.GBA) or a control GFP transgene (LVV.GFP) at similar number of vector integrations (LVV.GBA: 3.12 ± 0.15, LVV.GFP: 2.93 ± 0.14; Fig. 7a). As expected, the GCase activity in the cells transduced with the GBA transgene had significantly higher activity than those transduced with the GFP transgene (LVV.GBA: 3418.10% ± 612.2%, LVV.GFP: 20.38% ± 4.31%; Fig. 7b).

**Figure 7.**
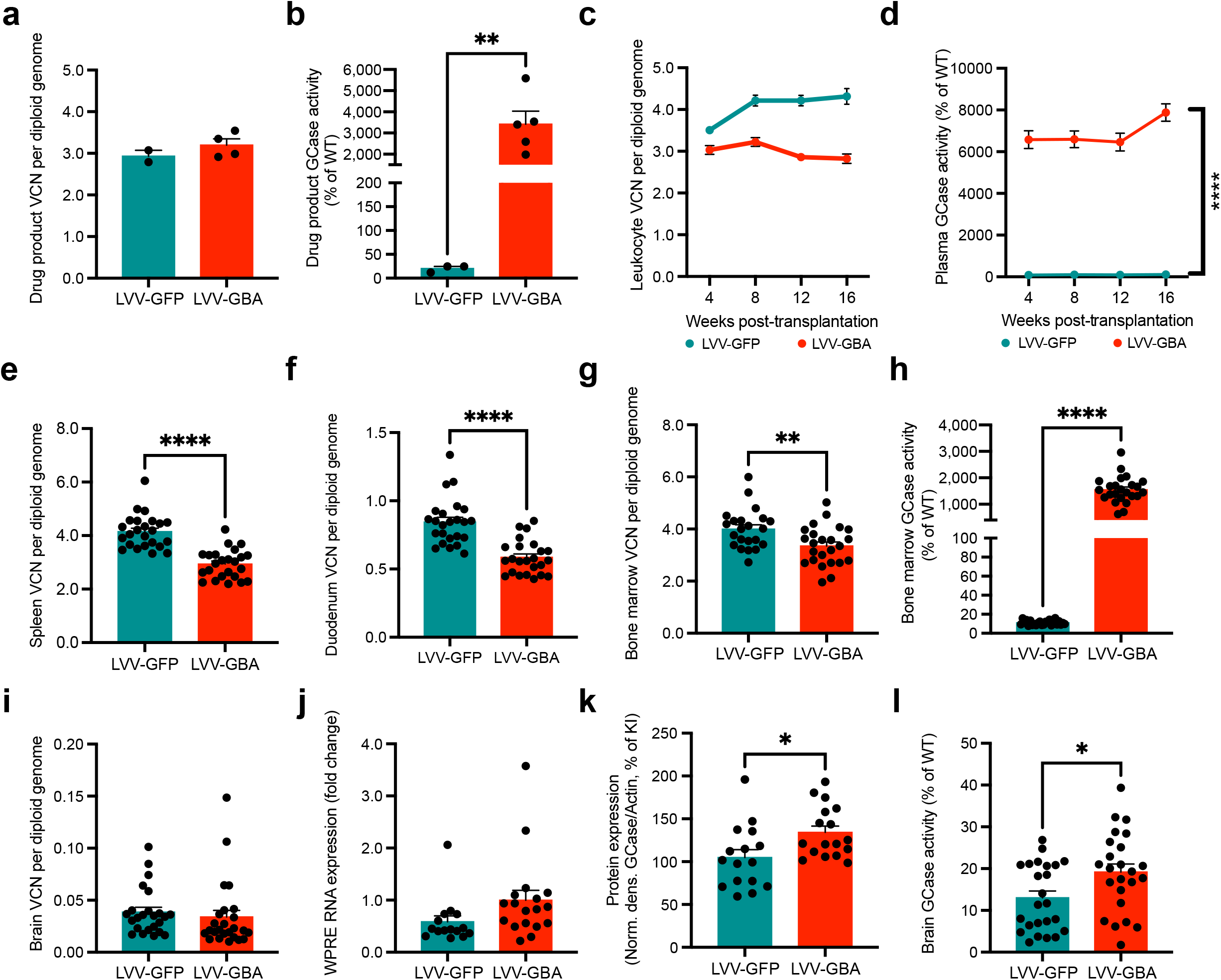
Biodistribution analysis after transplantation of LVV.GBA modified HSPCs into *Gba* mutant mice. **a** Vector copy number (VCN) in the drug product in the LVV-GFP and LVV-GBA treated kept in culture for four days in vitro. **b** GCase enzymatic activity in the drug product at DIV 4. **c-d** VCN and GCase enzymatic activity in white blood cells and plasma, respectively, measured longitudinally every 4 weeks post-transplantation. **e-f** VCN in spleen and duodenum at 16 weeks post-transplantation. **g-h** VCN and GCase enzymatic activity in the bone marrow at 16 weeks post-transplantation. **i-l** VCN, transgene RNA expression, GCase protein levels, and GCase enzymatic activity in the brain at 16 weeks post-transplantation. Bars represent means ± SEM. For a-b, e-l t-test was used for statistical analysis.

We then transplanted the genetically modified HSPCs into homozygous *Gba*^D409V^ knock-in mice, which have lower levels of enzyme activity in the bone marrow and brain compared to wild-type controls (Supp Fig 10a-d). Transplantation of LVV.GBA HSPCs led to detectable and stable levels of VCN in white blood cells (Fig. 7c). Furthermore, we observed GCase activity in the plasma starting at four weeks post-transplantation (earliest time point) till 16 weeks (terminal time point), whereas there is very minimal, if any, found in LVV.GFP-treated animals (Fig. 7d). At 16 weeks, we observed engraftment in the spleen (Fig. 7e), duodenum (Fig. 7f), and bone marrow (Fig. 7g) in both LVV.GBA-treated and LVV.GFP-treated animals. Importantly, we measured higher levels of GCase activity in the bone marrow of LVV.GBA-treated animals compared to animals transplanted with LVV.GFP HSPCs (LVV.GBA: 1547.75% ± 102.2%, LVV.GFP: 11.06% ± 0.50%; Fig. 7h). Within the brain we saw detectable levels of integrated vector (Fig. 7i) and RNA expression (Fig. 7j) in both animal groups. Moreover, we saw higher levels of GCase protein (LVV.GBA: 134.4% ± 7.24%, LVV.GFP: 105.0% ± 9.16%; Fig. 7k) and GCase enzymatic activity (LVV.GBA: 19.19% ± 1.93%, LVV.GFP: 13.00% ± 1.63%; Fig. 7l) in the LVV.GBA cohort versus the LVV.GFP cohort. Taken together, we observed widespread and robust biodistribution of the gene-modified cells in key organs, including the gastrointestinal tract, spleen, bone marrow, and central nervous system.

### HSPC-GT in a model of GRN-FTD increases progranulin and may address lysosomal dysfunction

We then applied our HSPC-GT platform to treat progranulin deficiency in a mouse model of genetic frontotemporal dementia (GRN-FTD). GRN-FTD is a familial form of neurodegeneration caused by the haploinsufficiency of GRN/progranulin, a lysosomal precursor protein that is widely expressed in the brain and secreted from microglia during frank disease^40^. Heterozygous and homozygous *Grn*^*R493X*^ mutant mice, a model of GRN-FTD^41^, were dosed with lineage negative cells transduced with high or low vector doses of LVV.GFP or LVV.GRN after standard (100 mg/kg) or increased (125 mg/kg) levels of busulfan conditioning. As in our previous studies, we observed biodistribution via vector integration quantification and transgene expression in the bone marrow and the brain after treatment for all treatment groups for LVV.GFP and LVV.GRN (Fig. 8a, b, e, f; Supp Fig. 11a-d). Across all LVV.GRN-treated groups, we measured supraphysiological levels of human progranulin via ELISA in the plasma and the bone marrow, equivalent to an average of 5-11-fold above previously reported wild-type levels (Fig. 8c, d)^42^. We also observed human progranulin levels in whole brain lysate of LVV.GRN-treated animals, equivalent to 30-50% of previously reported mouse progranulin levels in the brain (Fig. 8g)^43^. Using immunohistochemical visualization of mouse and human progranulin, we observed progranulin-positive cells in the cortex of LVV.GRN-treated homozygous mutant animals, consistent with the engraftment of LVV.GRN-positive cells in the brain (Fig 8i). Finally, and notably, increased level of busulfan conditioning resulted in significantly increased VCN, transgene expression, and GRN protein levels in the periphery and the brain (Fig. 8a-g).

**Figure 8.**
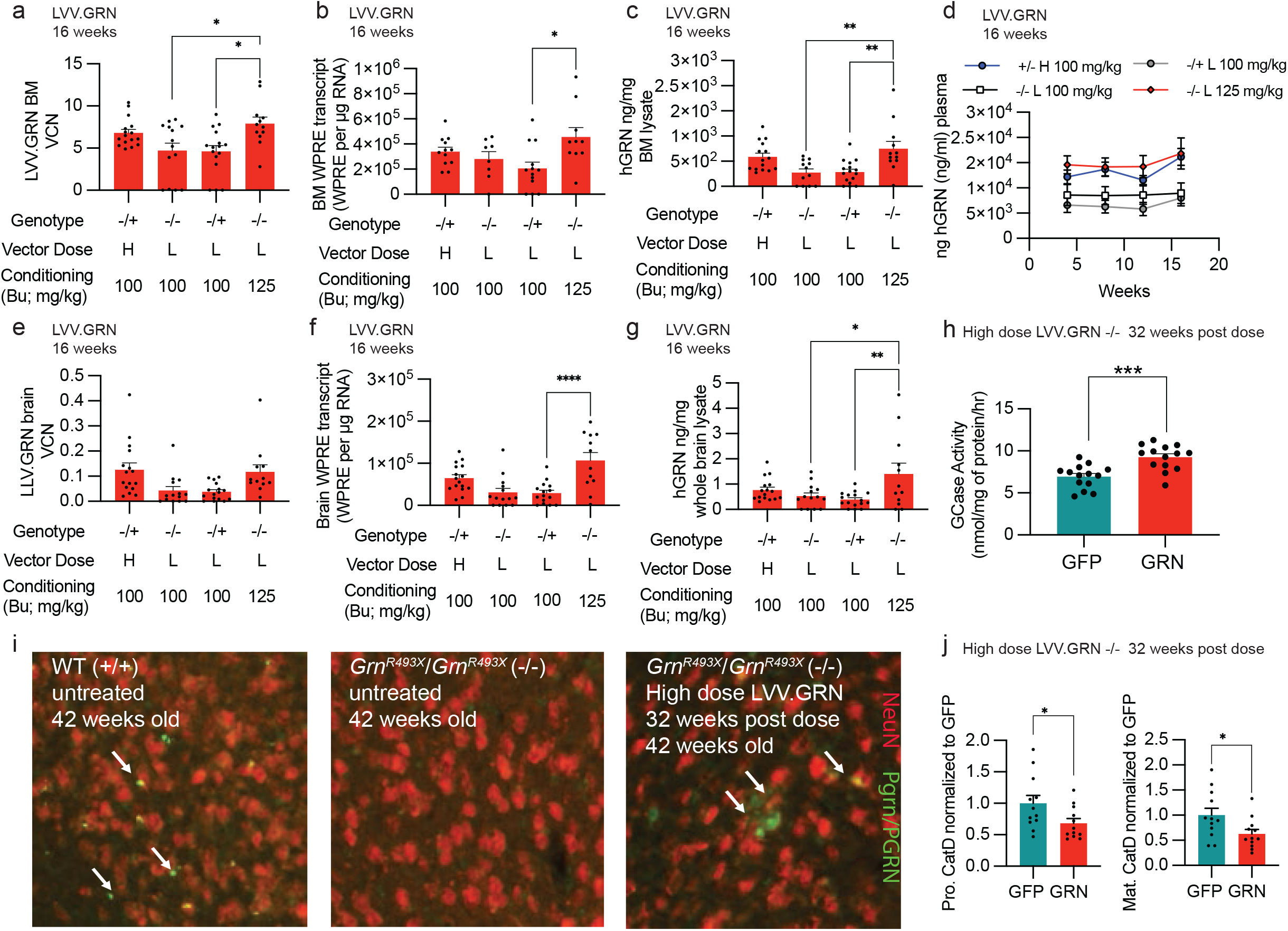
Genetically engineered HSPC administration in a mouse model of GRN-FTD shows widespread GRN in the periphery and the brain. **a-c** Vector copy number (VCN), transgene RNA, and human progranulin protein measurements in bone marrow for all LVV.GRN-treated groups at 16 weeks post-transplantation. **d** Longitudinal human progranulin protein measurements in plasma for all LVV.GRN-treated groups up to 16 weeks post-transplantation. **e-g** Vector copy number (VCN), transgene RNA, and human progranulin protein measurements in brain for all LVV.GRN-treated groups at 16 weeks post-transplantation. **h** GCase activity in whole brain lysate of homozygous mutant *Grn* mutant mice treated with LVV.GFP and LVV.GRN at 32 weeks post-transplantation. **i** Immunohistochemical imaging of neurons (NeuN, red) and progranulin (PGRN/Pgrn, green) in the cortex of untreated wild-type, homozygous mutant *Grn* mutant mice, or homozygous mutant *Grn* mutant mice treated with LVV.GRN. **j** Quantification of levels of the pro form and mature form of cathepsin D (CatD) in whole brain lysates of homozygous mutant *Grn* mutant mice treated with LVV.GFP and LVV.GRN at 32 weeks post-transplantation. Bars represent means ± SEM. Tukey’s multiple post-hoc comparison test was conducted for all statistical measures shown, with only significant differences (p<0.05) indicated.

While the reasons GRN deficiency leads to FTD are still unclear, progranulin is thought to play a role in regulating lysosomal homeostasis^44^. The protein and activity levels of several lysosomal enzymes, including decreased GCase and increased cathepsin D (CatD) have been observed in mouse models of GRN-FTD and human brain tissue from GRN-FTD patients^45,46^. Other models of GRN-targeted therapeutics have shown the ability to normalize these changes after treatment^45^. We observed similar normalization of lysosomal dysfunction after treatment, with LVV.GRN-treated homozygous *Grn* mutant animals 32 weeks post-transplantation showing increased GCase activity levels compared to LVV.GFP treated groups (Fig. 8h) and decreased mature cathepsin D protein (Fig. 8j) as compared to LVV.GFP treated groups.

## Discussion

We show here that HSPC-GT using an IV route of administration in mice results in the widespread engraftment of genetically engineered cells throughout the periphery and engraftment of MLCs in the brain. Similar levels of engraftment in the brain were previously reported by others^28,47^. In contrast, when HSPCs are administered via ICV, we observed engraftment is limited to the central nervous system. Strikingly, and in contrast to earlier work^18^, we observed significantly more engrafted MLCs using IV administration compared to ICV administration. While our methods and timepoints were similar for all key aspects, subtle technical disparities could explain the difference, like cell dose, injection location, and/or cell administration timing. Further studies will need to be conducted to explore these parameters and how they affect engraftment after ICV administration of HSPCs, if at all. We also observed that HSPC-derived MLCs were durably engrafted within the brain over a year post-transplantation and that these cells bore many of the transcriptional hallmarks of microglia, which strongly suggest they have the capacity to self-renew much like endogenous microglia. This is consistent with long-term follow-up of HSPC-GT in non-human primates, where HSPC-derived myeloid cells in the brain were observed ten years post-transplantation^15^. It is similarly consistent with the durable phenotypic rescue in most patients enrolled in clinical trials for MLD and CALD^48^. Importantly, these data strongly support the use of murine models for preclinical and translational research as they can effectively recapitulate the long-term engraftment dynamics of MLCs observed in humans.

Our comparison of MLCs to endogenous microglia suggests that most transcriptional features of myeloid cells in the brain are niche-dependent, as the two cell populations are largely similar. This observation parallels the ones from studies where monocytes and macrophages isolated from various tissues take on functional and transcriptional characteristics of tissue-resident macrophages of the lung, kidney, and liver when transplanted^49-52^. Furthermore, we show here that MLCs do not exhibit the signatures of invading macrophages associated with disease and injury, likely because these signatures are niche-dependent and reflect the damaged status of the brain. Our data shows, however, that MLCs and microglia can be clearly distinguishable by specific genes not reprogrammed during engraftment. Among these are *Apoe* and *Lyz2*, which are markers of Disease-Associated Microglia (DAMs)^53^. In the diseased brain where the DAM signature has been identified there is also significant recruitment of peripheral macrophages, which share similar cell surface markers to microglia^54^. One possibility is that the DAM signature could be in part driven by the developmental differences between bone marrow-derived myeloid cells and embryonic yolk sac-derived microglia, rather than by a disease state. How HSPC-derived MLCs and microglia differ on a functional level as a result of these differences in developmental origin will require further study.

Our work applying HSPC-GT in murine models of neurodegenerative disease shows that for both GBA-PD and GRN-FTD, engrafted cells have the capacity to produce protein at a potentially therapeutic level. Notably, gastrointestinal issues are a common symptom in Parkinson’s disease patients and mutations in *GRN* are implicated in cardiac dysfunction. A systemic gene therapy approach like HSPC-GT conceivably has advantages over other platforms including AAV gene therapies, where therapeutic effect is generally limited to either the periphery or the CNS. However, if CNS-restricted expression is therapeutically advantageous, our data suggest that ICV administration of genetically engineered HSPCs can achieve this.

Furthermore, previous work has demonstrated that AAV administration into the parenchyma leads to extremely high levels of transgene expression close to the injection site but a vastly lower expression level in distal brain regions^43,57^. Importantly, the cell type that expresses the therapeutic protein will depend on the AAV capsid and promoter used^58^. Conversely, we showed here that the tiling behavior of MLCs expands the reach of HSPC-derived cells to the whole brain, resulting in an even and widespread distribution of genetically modified cells. It is worth noting that AAV-mediated delivery of GBA and GRN to neurons in mouse models of GBA-PD and GRN-FTD have been successful in modifying disease-associated phenotypes^45,59^. Our biodistribution studies and our initial biochemical characterization of lysosomal dysfunction in *Grn* mutant mice support the capacity of reaching similar results with HSPC-GT. Further characterization is required to determine whether HSPC-derived MLCs can provide the levels and modality of therapeutic transgene expression necessary to correct the wider functional changes in the brain associated with these disorders.

While this work addresses fundamental questions around HSPC-GT in the CNS, such as the identity of MLCs relative to endogenous microglia and the durability of MLC engraftment, there are broader questions that remain unresolved. These include understanding how the myeloablative agent acts in the CNS. Our results clearly demonstrate busulfan conditioning is instrumental in enabling brain engraftment with higher doses leading to higher engraftment and protein expression. Additionally, the identity of the cell that crosses the blood brain barrier (e.g., HSPC, myeloid progenitor cell) after intravenous administration or that first engrafts after ICV administration is unclear. It also worth examining if injection of myeloid cells alone could achieve similar levels of engraftment. Moreover, future studies should assess what the window of opportunity is for cells to cross the blood-brain barrier following myeloablation.

In conclusion, our work provides critical information substantially expanding our understanding of the nature of microglia-like cells derived from long-term HSPCs and opens new avenues for the potential exploitation of genetically engineered MLCs for the treatment of common and devastating neurodegenerative disorders such as frontotemporal dementia and Parkinson’s disease.

## Methods

### Mouse models and tissue collection

Comparisons of intravenous (IV) and intracerebroventricular (ICV) routes of administration were completed using wild-type (WT) C57BL/6J mice (Jackson Laboratory, stock number 000664). Proof of concept experiments for GBA-Parkinson’s Disease (GBA-PD) were completed using homozygous *Gba*^*D409V/D409V*^ mutant mice (Jackson Laboratory, C57BL/6N-*Gba*^*tm1.1Mjff*^/J, stock number 019106)^39^ and wild-type C57BL/6NJ mice (Jackson Laboratory, stock number 005304) as controls. Proof of concept experiments for progranulin-associated frontotemporal dementia (GRN-FTD) were completed using heterozygous and homozygous *Grn*^*R493X*^ mutant mice on the C57BL/6J background^41^. Donor and recipient animals for all experiments were 8-12 weeks of age. Terminal collection for all animals occurred 14-16 weeks after cell administration except for the long-term cohort for scRNA-Seq experiments (12-13 months post-transplantation) and the long-term cohort for the GRN-FTD proof of concept study (32-36 weeks post-transplantation). At necropsy, animals first underwent whole-body transcardial perfusion either heparinized saline or PBS followed by tissue harvesting. Samples for biochemistry analysis were flash frozen and stored at -80°C. Samples for IHC analysis were fixed in 4% PFA in PBS overnight at 4°C.

### Generation of genetically engineered HSPC drug product

For all studies, lineage negative cells were enriched from total bone marrow isolated from femurs and tibias of donor mice (6-12 weeks of age) using the EasySep Mouse Hematopoietic Progenitor Cell Isolation Kit (STEMCELL Technologies) in conjunction with the RoboSep-S (STEMCELL Technologies) automated cell isolation machine and confirmed by flow analysis. Cells were stained for lineage markers with a PE-Cy5-conjugated lineage marker cocktail including B220, TER119, TCR-β, CD8a, CD3e, CD4, Ly6G/Ly6C, and CD11b, and with hematopoietic stem cell markers APC-conjugated anti-CD117 (c-Kit) and PE-conjugated Ly6A/E (Sca-1) (BD Biosciences). Lineage negative cells were transduced with a lentiviral vector in a cell incubator at 2 ×10^6^ cells/mL in either StemSpan SFEM II (STEMCELL Technologies) or StemMACS HSC Expansion Media (Miltenyi Biotec) growth media freshly supplemented with TPO (10 ng/mL), SCF (100 ng/mL), and FLT3 (50 ng/mL). Approximately 16-22 hours later, cells were collected, washed at least three times with DPBS (without Ca^2+^ and Mg^2+^), and resuspended in an appropriate volume for dosing. A small number of cells (3,000 to 4,000 cells) were set aside for use in the colony forming unit assay using M3434 media (STEMCELL Technologies) and analyzed 7-10 days later.

### Conditioning of recipient animals and drug product administration

Four days prior to cell administration, recipient animals (6-12 weeks of age) received daily intraperitoneal injections of busulfan (Busulfex, Otsuka Pharmaceutical) at 25 mg/kg for a cumulative dose of 100 mg/kg or 125 mg/kg. For comparisons of routes of administration, drug product pools were generated from equal number of male and female mice. For single-cell RNA-Seq experiments, drug product pools were generated from male mice only. For both IV and ICV administration animals were dosed with 5×10^5^ cells. For animals dosed using both IV and ICV, animals were dosed with 5×10^5^ for both routes for a total of 1×10^6^ cells. Five days after drug product administration, all animals despite route of administration received an additional IV administration of non-transduced total bone marrow cells (5×10^5^ cells in 50 μL per animal, resuspended in DPBS without Ca^2+^ or Mg^2+^) to support the survival of animals that only received ICV administration of drug product. Subsequent studies (GBA-PD and GRN-FTD proof of concept studies) only administered drug product intravenously and the second total bone marrow administration was eliminated. Additionally, in these studies, donors and recipients were sex matched and for the GRN-FTD proof of concept study an additional analysis of five days of busulfan conditioning was added.

### Lentiviral vector generation

VSV-G pseudotyped third generation self-inactivating lentiviral vectors for LVV.GFP, LVV.GRN, and LVV.GBA were commercially generated (University of Cincinnati Vector Production Facility, VIVEbiotech, VectorBuilder) using HEK293T cells and standard protocols. Lentiviral vector functional titers ranged from 1.7×10^8^ to 3×10^9^ TU/mL.

### Flow cytometry analysis for myeloid and lymphoid populations

For the blood and bone marrow samples, red blood cells were lysed and resuspended in RPMI medium. White blood cells were split into aliquots. The first aliquot was stained with CD45.1-APC (clone A20), CD45.2-PE (clone 104), CD3ε-PerCP-Cy5.5 (clone 17A2), and B220-APC-Cy7 (clone RA3-6B2) for 20 minutes at 4°C. The second aliquot was stained with CD45.1-APC (clone RB6-8C5), CD45.2-PE (clone 104), TER119-PerCP-Cy5.5, CD11b-PE-Cy7 (M1/70), GR1-APC-Cy7 (clone RB6-8C5) for 20 minutes at 4°C. After two washes, both aliquots were incubated with the viability marker Sytox Blue for 8 minutes at room temperature. Stained cells were measured using a MACSQuant Analyzer 10 (Miltenyi Biotec) and analyzed using FlowJo (FlowJo, LLC).

### Quantification of GFP and Iba1 positive cells in the brain

Five sections across the rostral caudal axis of the brain (two sections from approximately +0.2 mm to +3.2 mm relative to bregma, two sections from approximately +0.2 mm to -4.06 mm relative to bregma, and one section covering the cerebellum from approximately -5.2 mm to -6.1 mm to bregma) were uploaded to the HALO platform (Indica Labs, version 3.1.1076.301) for analysis. A tissue classifying algorithm was trained to detect and select tissue from background, thereby creating a region of interest for subsequent analysis. Analysis outputs for each image set included positive signal area for both stains, colocalized positive signal area, cell count for each stain, and colocalized cell count.

### SDS-PAGE of beta glucocerebrosidase, progranulin, and Cathepsin D

For RAW264.7 cells, 2.5 μg of protein lysate was mixed with 4x protein sample buffer (XT Sample Buffer, Bio-Rad) and water to a volume of 15 μL. The samples were heated at 95°C for 5 minutes. Samples were loaded into precast 10% Bis-Tris polyacrylamide gels (Bio-Rad) with MOPS buffer (Bio-Rad) and electrophoresed. For brain lysates, 10 μg of protein lysate was mixed with 4x protein sample buffer (Invitrogen or Bio-Rad) and water or lysis buffer to a volume of 15 μL. The samples were heated at 70-95°C for 5-10 minutes. Samples were loaded into precast 4-20% Tris-HCl polyacrylamide gels (Bio-Rad) with Tris/Glycine/SDS buffer (Bio-Rad) and electrophoresed. After completion of electrophoresis, gels were transferred to PVDF membranes using the iBlot 2 system (ThermoFisher). PVDF membranes were blocked for 1 hour with 5% BSA in 0.05% PBS-Tween. Primary antibodies (GCase, progranulin, cathepsin D, beta-actin) were diluted in 5% BSA in 0.05% PBS-Tween and incubated with PVDF membranes overnight at 4°C. PVDF membranes were washed three times with 0.05% PBS-Tween and then incubated with appropriate HRP secondary antibodies (Bio-Rad) for 1 hour at 1:50,000 dilution in 5% BSA in 0.05% PBS-Tween. Blots were washed three times in 0.05% PBS-Tween for 10 minutes each and then incubated with chemiluminescence reagent (SuperSignal West Femto, ThermoFisher). Images were acquired using a charge-coupled camera (G:BOX, Syngene). Bands were quantified using ImageJ.

### Transduction of immortalized cell lines

RAW264.7 mouse macrophage cells (ATCC; confirmed by STR profiling, IDEXX) were transduced with increasing multiplicity of infections with LVV.GRN or LVV.GBA vector. Approximately 85,000 cells per well of a 6-well dish were seeded 24 hours prior to transduction in 2 mL of complete growth media (DMEM supplemented with 10% heat-inactivated FBS, pyruvate, GlutaMAX, and penicillin/streptomycin). Transduction media was removed after 16 hours and replaced with fresh complete growth media. Cells were collected 4 days post-transduction, washed with DPBS twice, pelleted by centrifugation at 500xg for 5 min, and frozen at -80°C prior to downstream molecular characterization.

### Sample preparation and molecular characterization

Genomic DNA (gDNA) was isolated using the QIAamp DNA Mini QIAcube Kit (Qiagen) in conjunction with the QIAcube (Qiagen). Immortalized cell line gDNA was isolated using the DNeasy Blood & Tissue kit (Qiagen). Vector copy number (VCN) quantification was completed either by digital droplet PCR (ddPCR) or quantitative PCR (qPCR). Specific primers targeting the WPRE element (WPRE v1) or HIV Psi element were used to detect the integrated lentiviral vector and specific primers targeting *Gtdc1* or *Tfrc* were used as genomic reference (Supp. Table 1).

RNA isolation and reverse transcription for murine samples from the GRN-FTD proof of concept study was completed using the QIAsymphony RNA kit (Qiagen). Absolute quantification of transgene copies per microgram of tissue was completed using a standard curve of in vitro transcribed WPRE RNA and specific primers and probes targeting WPRE (WPRE v2, Supp. Table 1). RNA isolation and reverse transcription for all other samples was completed using the PureLink RNA Mini kit (ThermoFisher) followed by SuperScript IV VILO with ezDNase (ThermoFisher). Specific primers and probes targeting the WPRE element (WPRE v1; Supp. Table 1) of the integrating lentiviral vector and *Actb* (ThermoFisher), as a housekeeper, were used. Primers and probes, cDNA, TaqMan Fast Advanced Master Mix (ThermoFisher) were combined according to the manufacturer’s guidelines and amplified using a QuantStudio 7 Flex (ThermoFisher). The 2^-ΔΔCt^ method was used to calculate transgene expression.

Protein was extracted using a detergent-based lysis buffer dependent on the assay followed by removal of cellular debris by centrifugation at 14,000xg for 15 minutes. Protein concentration was determined using the Pierce BCA Protein Assay Kit (ThermoFisher). SDS-PAGE was used to measure beta glucocerebrosidase (GCase) and progranulin protein expression. GCase enzymatic activity was measured using the artificial substrate 4-methylumbelliferyl β-D-glucopyranoside (4-MUG) following a similar method to previously published^60,61^. Progranulin protein levels were measured using a human-specific ELISA (Quantikine and DuoSet; R&D Systems) according to the manufacturer’s instructions, with minor modifications for the measurement of human progranulin in brain lysates. For brain samples, reagents were diluted in 0.05% Tween 20 in Tris-buffered saline (TBS). 75 μg of protein was loaded of mouse brain lysate and 1.5ug of protein was loaded for mouse bone marrow, and plasma was measured at a final dilution of 1:500 for the human progranulin ELISA assay. 30 to 500 ng of protein was used to measure transgene expression of progranulin in RAW264.7 cells.

### Immunofluorescence staining of mouse

For quantification of engraftment within the central nervous system, brains were paraffin embedded, sectioned at 20 μm, and 12 sections were stained for GFP and Iba1 and 12 sections were stained for GFP and TMEM119 at a 300 μm interval. Slides were scanned using VS200 Research Slide Scanner (Olympus). Images were uploaded and analyzed using the HALO platform (v3.1.1076.301, Indica Labs).

For qualitative analysis of GFP engraftment, peripheral tissues were fixed overnight in 4% PFA and then transferred to a solution of 30% sucrose until they sank to the bottom. Subsequently, tissues were mounted in tissue freezing media and stored at -80°C. Tissues were sectioned at 20-50 μm (depending on tissue) using a sliding microtome with a frozen stage (ThermoFisher) or cryostat (Leica). Sections were mounted on glass slides using a glycerol-based mounting media.

For progranulin IHC, brains were processed similar to peripheral tissues. Sections were first incubated with an antigen retrieval solution (Antigen Retrieval Reagent-Basic, R&D Systems) at 80°C for 30 minutes followed by blocking (2% non-fat dry milk, 0.3% Triton X-100, and 0.01% sodium azide in PBS). Next, sections were blocked again in 1% donkey serum in 0.3% Triton X-100 in PBS for 1hr at room temperature. Sections were then incubated with primary antibodies for progranulin (AF2420 R&D Systems) and NeuN (MAB377 MilliporeSigma) overnight at 4°C followed by the appropriate fluorescent secondary antibodies for one hour.

### Isolation of microglia/MLCs and single cell RNA-Seq

After transcardial perfusion, the brains of one male and female mice from IV and ICV groups were processed by enzymatic digestion (Neural Tissue Dissociation Kit (P), Miltenyi Biotec). The resulting single cell suspension was stained with viability dye and antibodies for microglia cells, neurons, astrocytes and endothelial cells (LIVE/DEAD Fixable Aqua Dead Cell Stain Kit, ThermoFisher; CD45.1 clone A20 in BV421 (BD); CD45.2 clone 104 in BV421 (BD); CD11b clone M1/70 in APC-780 (ThermoFisher); CX3CR1 clone SA011F11 in BV605 (BioLegend); PECy7 Thy1.1 (also known as CD90.1) clone OX-7 in PE-Cy7 (BD); Thy1.2 (also known as CD90.2) clone 53-2.1 in PE-Cy7 (BD); ACSA-2 clone IH3-18A3 in PE (Miltenyi Biotec); CD31 clone 390 in PerCP/Cy5.5 (BioLegend). GFP-positve and GFP-negative fractions of microglia cells, defined as CD45+ CD11b+ Cx3cr1+, were FACS-sorted using a MA900 Multi-Application Cell Sorter (Sony Biotechnology). For the generation of single cell transcriptomes, 5×10^3^ target cells from each sorted population were run using the Chromium Controller (10x Genomics) using Chromium Next GEM Single Cell 3’ Reagent Kits (10x Genomics). The libraries generated were then run on the NextSeq 550 Sequencing System (Illumina) using NextSeq 500/550 High Output v2.5 (150 cycles) Kit (Illumina). The Illumina raw BCL sequencing files were processed through the CellRanger software (10x Genomics) for generating FASTQ files and count matrixes (https://support.10xgenomics.com/single-cell-gene-expression/software/overview/welcome). The count matrixes were then used as input for the SEURAT V4.0 (https://satijalab.org/seurat/) and MONOCLE3 (http://cole-trapnell-lab.github.io/monocle-release/) R tools for single cell genomics analyses. Briefly, single cell barcodes were filtered for the ones containing mitochondrial gene content lower than 15%. Expression data then were normalized, scaled, and searched for variable features using the *SCTransform* function of SEURAT V4.0 followed by tSNE dimensionality reduction and clustering using the *FindClusters* function with resolution set at 0.2. For generating the maps shown in Fig.3b and Fig.4a we run the tSNE coordinates on the R package *ggplot2* (https://cran.r-project.org/web/packages/ggplot2/index.html). For visualization of gene expression shown in Fig.2b the SEURAT object was converted into a MONOCLE object and individual genes were plotted using the *plot_cells* function of MONOCLE3. The heatmaps of Fig. 3e, 4d and 5a were generated based on the average normalized gene expression per sample using the R package pheatmap (https://cran.r-project.org/web/packages/pheatmap/index.html).

### Statistics

Statistical analysis was performed using either Prism (GraphPad) or R. For analyzing the relationship between vector volume and transgene metrics in Fig.6, we fit a linear regression model and measured the Pearson correlation and its statistical relevance (p-value). For scRNA-Seq data we used GSEAPreranked analysis to identify the gene sets that are enriched in GFP-negative and GFP-positive samples. To perform this analysis, we firstly derived a score using log2FoldChange and p-value from differential gene expression analysis (score=-s*log10(p-value) with s=-1 if the log2FoldChange of gene <0, otherwise s=+1). We then sorted the genes based on the score value in decreasing order and used the whole ranked gene list file (including 11,992 genes and their score values) as input for the online tool WebGestalt (http://www.webgestalt.org) to perform GSEAPreranked analysis. We then generated a network-based enrichment map visualization for the 20 enriched gene sets. In this network, the edges correspond to the overlapping between connected gene sets and were calculated using the gene list of the leading-edge genes of each gene set from GSEA. The node size instead indicates the number of the leading-edge genes. The overlapping score was calculated using the following formula: 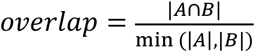 where |*A* ∩ *B*| is the cardinality of the intersection between gene sets A and B while |*A*| and |*B*| are the cardinality of gene set A and gene set B, respectively. For the plot shown in Fig. 3D we applied the Procrustes tool from the R Package Vegan (https://cran.r-project.org/web/packages/vegan/vegan.pdf) to perform unsupervised clustering on the global gene*cell expression matrixes in different sample types. Since the number of cells for sample types are different, we downsampled all sample types to the smallest dataset and then calculated Procrustes distance for each sampled data sets and calculated and plotted the average from 10 random samplings. We then perform clustering using the averaged Procrustes distance. For all other data, a Student t-test (when only two groups were present), one-way ANOVA with Tukey’s post hoc analysis (when multiple groups were present), or linear regression was used. Significance was defined as p < 0.05.

## Acknowledgements

We thank all the members of AVROBIO, Inc. for their continued support of our work. In particular, we would like to thank Maurine Braun, Maria Grigorova, Christine Oborski, Daniella Pizzurro, Jurgen Poci, Steven Tyler, Claudia Harper, Carolina Romano, Steven Avruch, Monique da Silva, Geoff MacKay, and Deanna Petersen for scientific support and assistance. We would like to also thank Taneli Heikkinen, Teija Parkkari, Sarah Davis, Karen Wong, Daphne Gordon, and Jaime Nederhoed at Charles River Laboratories (CRL) for study execution and diligent care of study animals. Additionally, we thank Scott W. Allen (BioAgilytix), Kelly Colletti (CRL), Katherine Domingue (QPS), Romain Genard (CRL), Lauriane Padet (CRL), Kaisa Paldanius (CRL), Taina-Kaisa Stenius (CRL), and Jenifer Vija (CRL) for their contributions to sample analysis. The *Grn*^R493X^ mice were generated in the Robert Farese Laboratory at The J. David Gladstone Institutes with grant funding provided by the Consortium for Frontotemporal Dementia Research and under a subaward of a University of California, San Francisco (UCSF) grant from the Alzheimer’s Disease Center (ARDC) of the National Institutes of Health. The work was funded by AVROBIO, Inc. Certain schematic elements were generated using BioRender.com.

## Author Contributions

RNP and MPD contributed equally to the design, execution, and reporting of all work presented. ML performed FACS-sorting of cell subsets and generated transcriptome libraries. CB contributed to setup and execution of scRNA-Seq library preparation. YD and ZU contributed to the execution of in vivo studies. JWS and JKY contributed to execution and reporting of cell sorting assays. LB contributed to execution of VCN assay and reporting. FH, QN, and TP contributed to the execution of molecular assays (ELISA, VCN, Western Blots, and Colony Forming Unit Assay). OC contributed to the initial designs of all studies presented and the execution of the biodistribution experiments. NPT designed the lentiviral vectors. AY and LB contributed to the statistical analysis and reporting of the scRNA-Seq data. CM contributed to the design and the reporting of all work presented.

