## Supplemental Materials for "Genetically engineered microglia-like cells have therapeutic potential for neurodegenerative disease"

### Supplemental Table 1. Primers and probe sequences

| Target | Purpose | Primer | Sequence (5'-3') | Vendor |
|---|---|---|---|---|
| WPRE (v1) | VCN, Expression | Forward | TTCTGGGACTTTCGCTTTCC | IDT |
| WPRE (v1) | VCN, Expression | Reverse | CCGACAACACCACGGAATTA | IDT |
| WPRE (v1) | VCN, Expression | Probe | ATCGCCACGGCAGAACTCATCG | IDT |
| *Tfrc* | VCN | Forward | Not available | ThermoFisher, 4458367 |
| *Tfrc* | VCN | Reverse | Not available | ThermoFisher, 4458367 |
| *Tfrc* | VCN | Probe | Not available | ThermoFisher, 4458367 |
| *Gtdc1* | VCN | Forward | GAAGTTCAGGTTAATTAGCTGCTG | IDT |
| *Gtdc1* | VCN | Reverse | GGCACCTTAACATTTGGTTCTG | IDT |
| *Gtdc1* | VCN | Probe | ACGAACTTCTTGGAGTTGTTTGCT | IDT |
| *Actb* | Expression | Forward | Not available | ThermoFisher, Mm02619580_g1 |
| *Actb* | Expression | Reverse | Not available | ThermoFisher, Mm02619580_g1 |
| *Actb* | Expression | Probe | Not available | ThermoFisher, Mm02619580_g1 |
| WPRE (v2) | Expression | Forward | GGCTTTCRTTTTCTCCTCCTTGTAT | ThermoFisher |
| WPRE (v2) | Expression | Reverse | CGGGCCACAACTCCTCATAA | ThermoFisher |
| WPRE (v2) | Expression | Probe | AATCCTGGTTGCTGTCTC | ThermoFisher |
| Psi element | VCN | Forward | CAGGACTCGGCTTGCTGAAG | IDT |
| Psi element | VCN | Reverse | TCCCCCGCTTAATACTGACG | IDT |
| Psi element | VCN | Probe | CGCACGGCAAGAGGCGAGG | IDT |

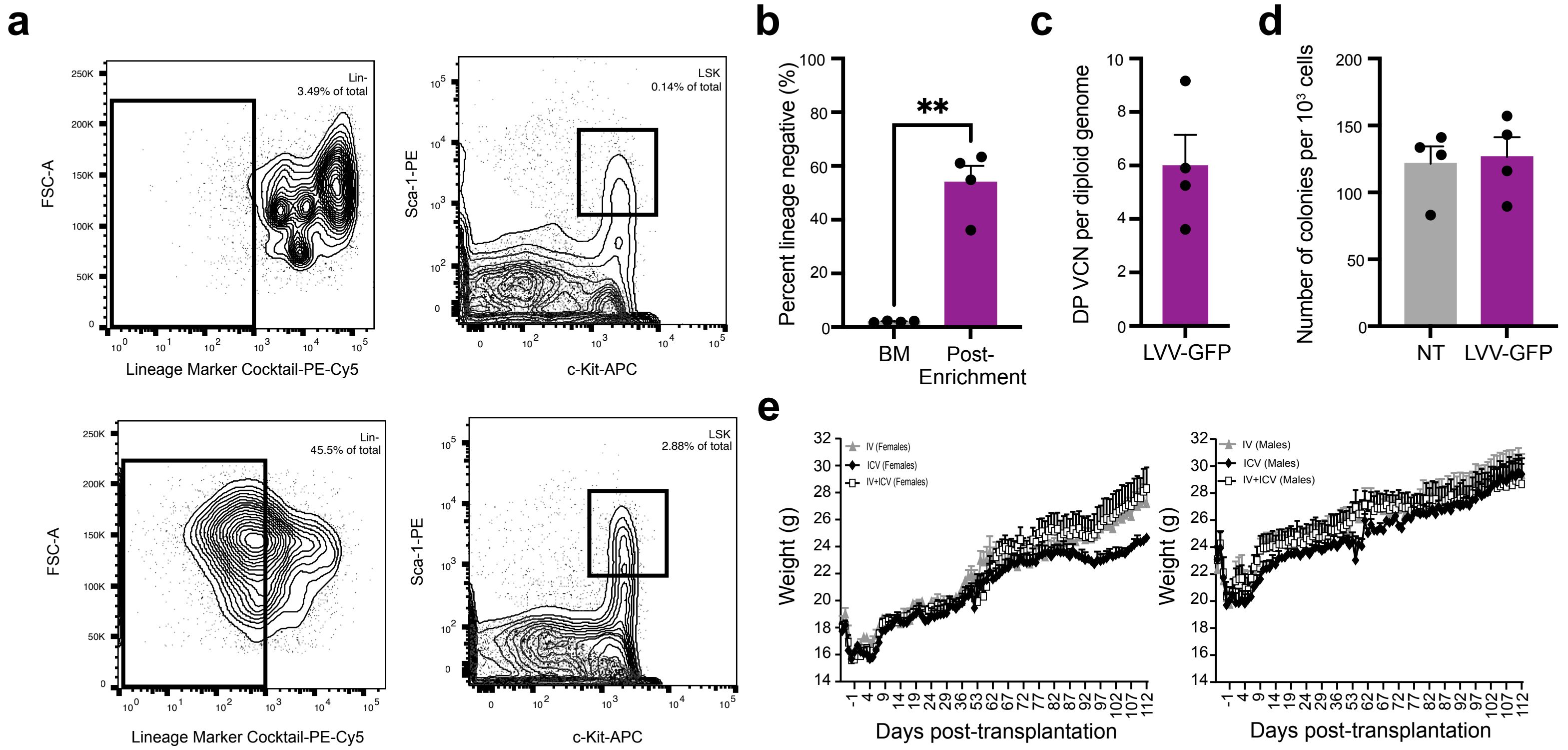


**Supplementary Figure 1. Drug product characterization and post-transplantation body weights.**
**a** Representative flow cytometry analysis plots of total bone marrow (top) and lineage negative cells after enrichment (bottom) for lineage markers (left panels) and c-Kit/Sca1 (right panels). **b** Quantification of lineage negative population in four drug product prepraration s. **c** Vector copy number per diploid genome in the drug product. **d** Colony forming unit assay to assess pluripotency of lineage negative cells. **e** Body weights of female (left) and male (right) animals through the course of the study. T-test was used for statistical analysis. **, p<0.01. Bars represent means ± SEM.

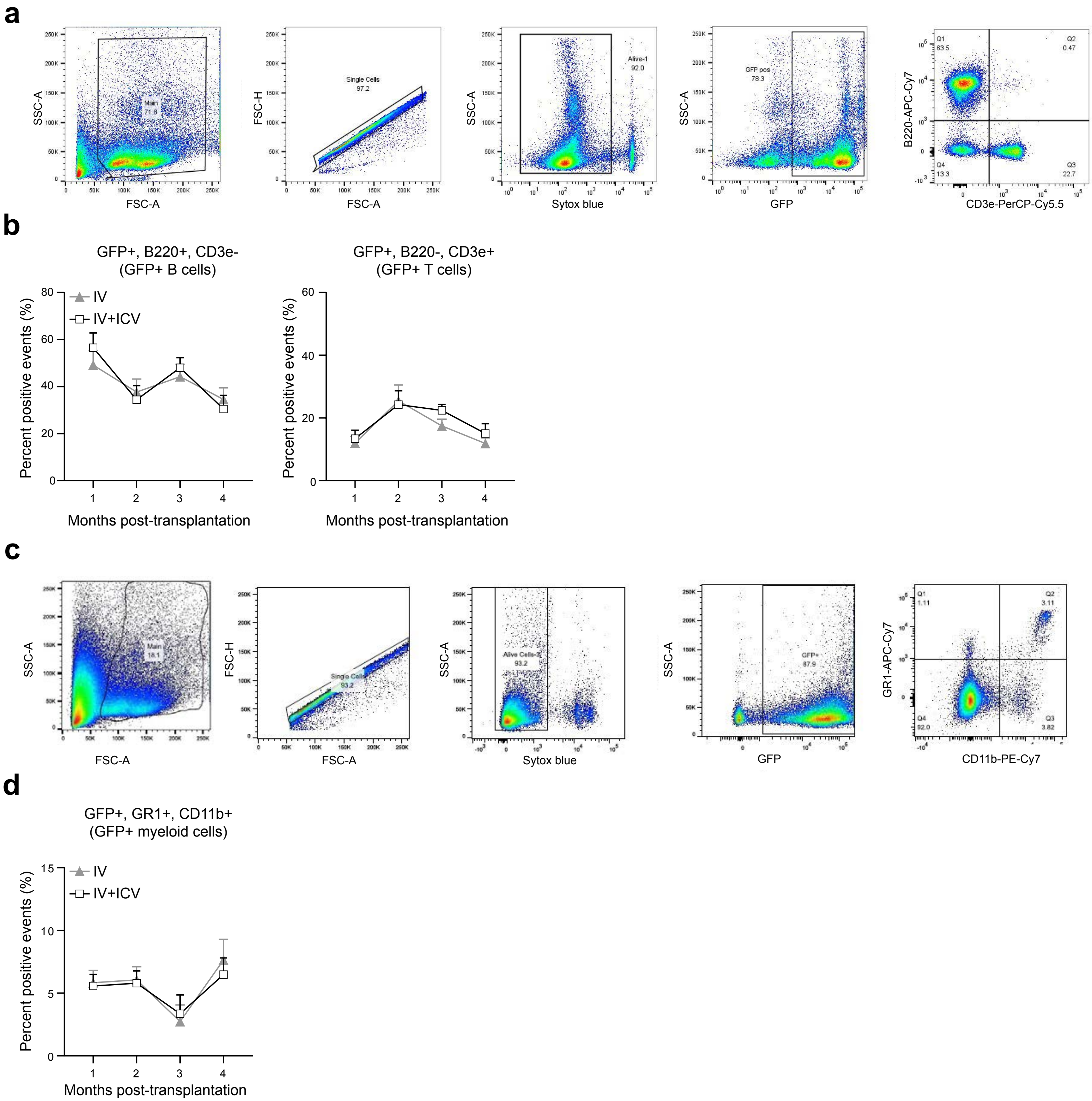


**Supplementary Figure 2. Flow cytometry analysis of peripheral blood to measure engraftment in various cell compartments. a-b** Representative flow cytometry plots and quantitation of GFP-positive (GFP+) B cells (B220+, CD3e-) and T cells (B220-, CD3e+) in IV and IV+ICV animals. **c-d** Representative flow cytometry plots and quantitation of GFP-positive myeloid cells (GR1+, CD11b+) in IV and IV+ICV animals. Symbols represent means ± SEM.

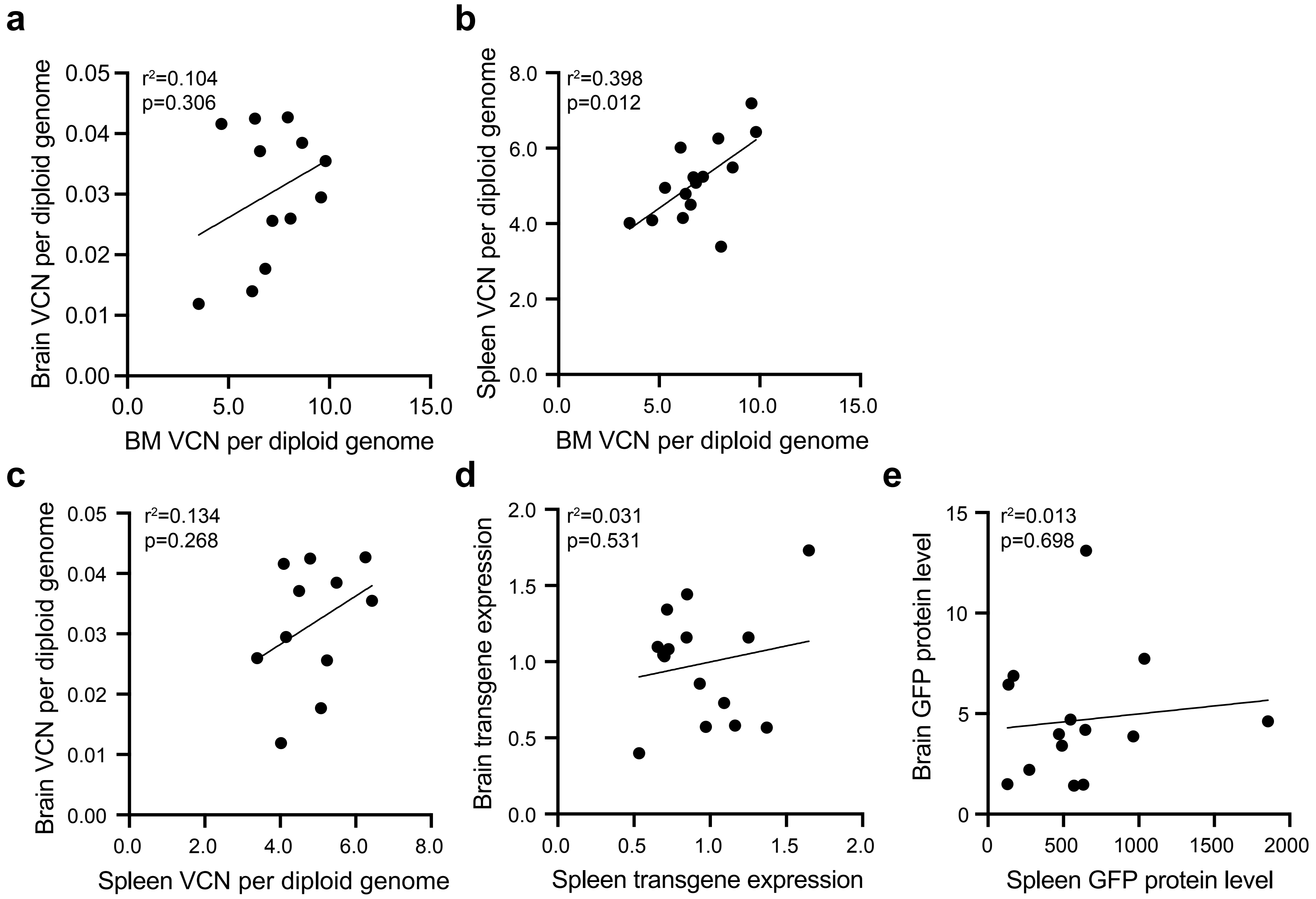


**Supplementary Figure 3. Linear regression analysis of brain, bone marrow, and spleen biodistribution metrics from animals dosed IV or IV+ICV.**

**a** Brain vector copy number (VCN) versus bone marrow (BM) VCN per genome. **b** Spleen VCN versus BM VCN. **c** Brain VCN versus spleen VCN. **d** Brain RNA vs spleen transgene RNA**. e** Brain GFP protein versus spleen GFP protein. Line and statistics represent linear fit after simple linear regression.

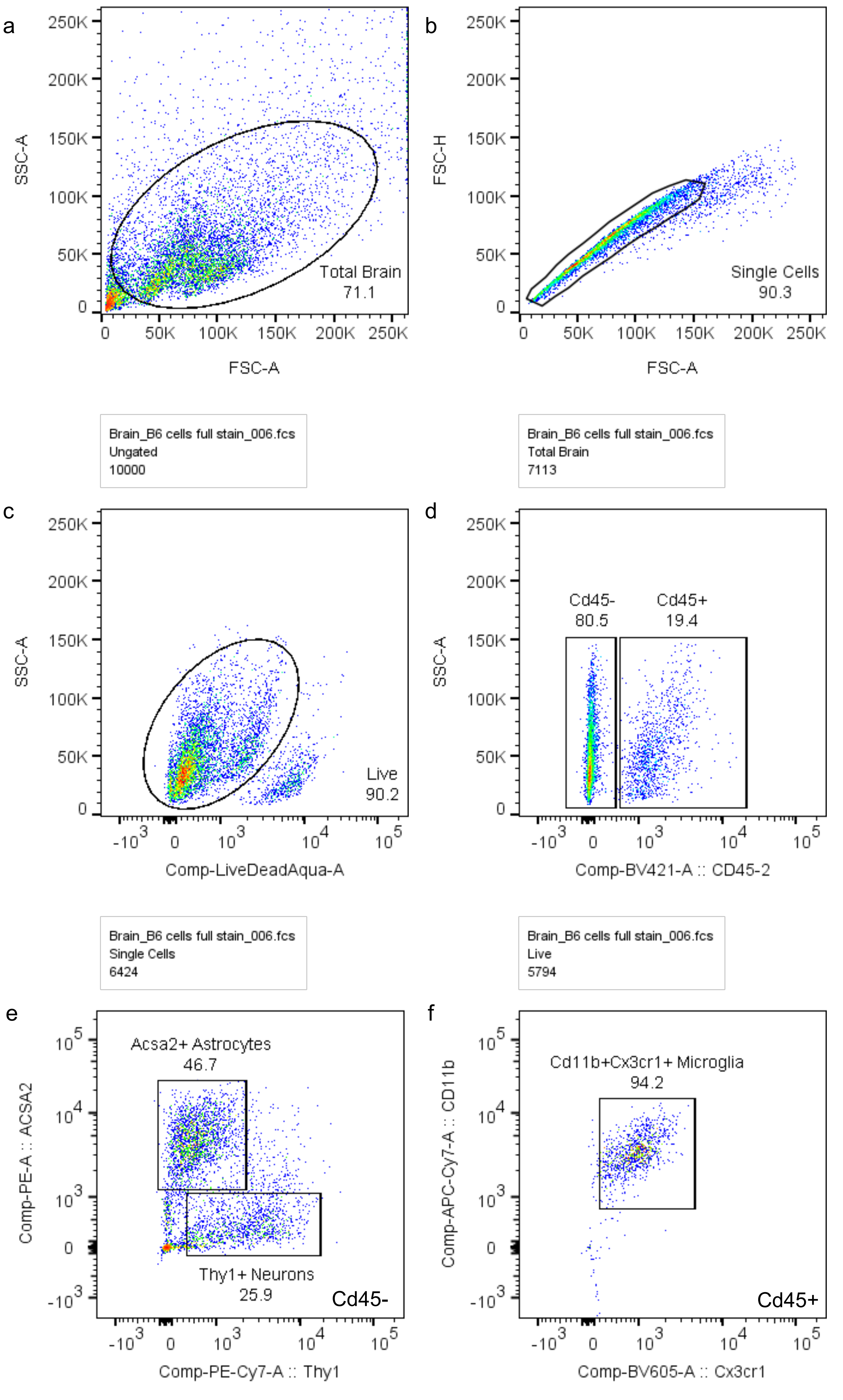


**Supplementary Figure 4. Gating strategy for FACS isolation of microglia, astrocytes, and neurons**. Whole adult mouse brains were enzymatically digested to single cell suspension and then stained with cell-surface marker antibodies to enable FACS-assisted isolation of specific cell populations. Representative gating strategy shows the selection of the single cell population (**a**,**b**), selection for viability (**c**), gating on the immune cell marker CD45 (**d**) selection of CD45- ACSA2+, Thy1- astrocytes, and CD45- ACSA2- Thy1+ neurons (**e**), selection of CD45+ CD11b+, C3X3CR1+ microglia (**f**). Percentages of parent population are listed next to each gate.

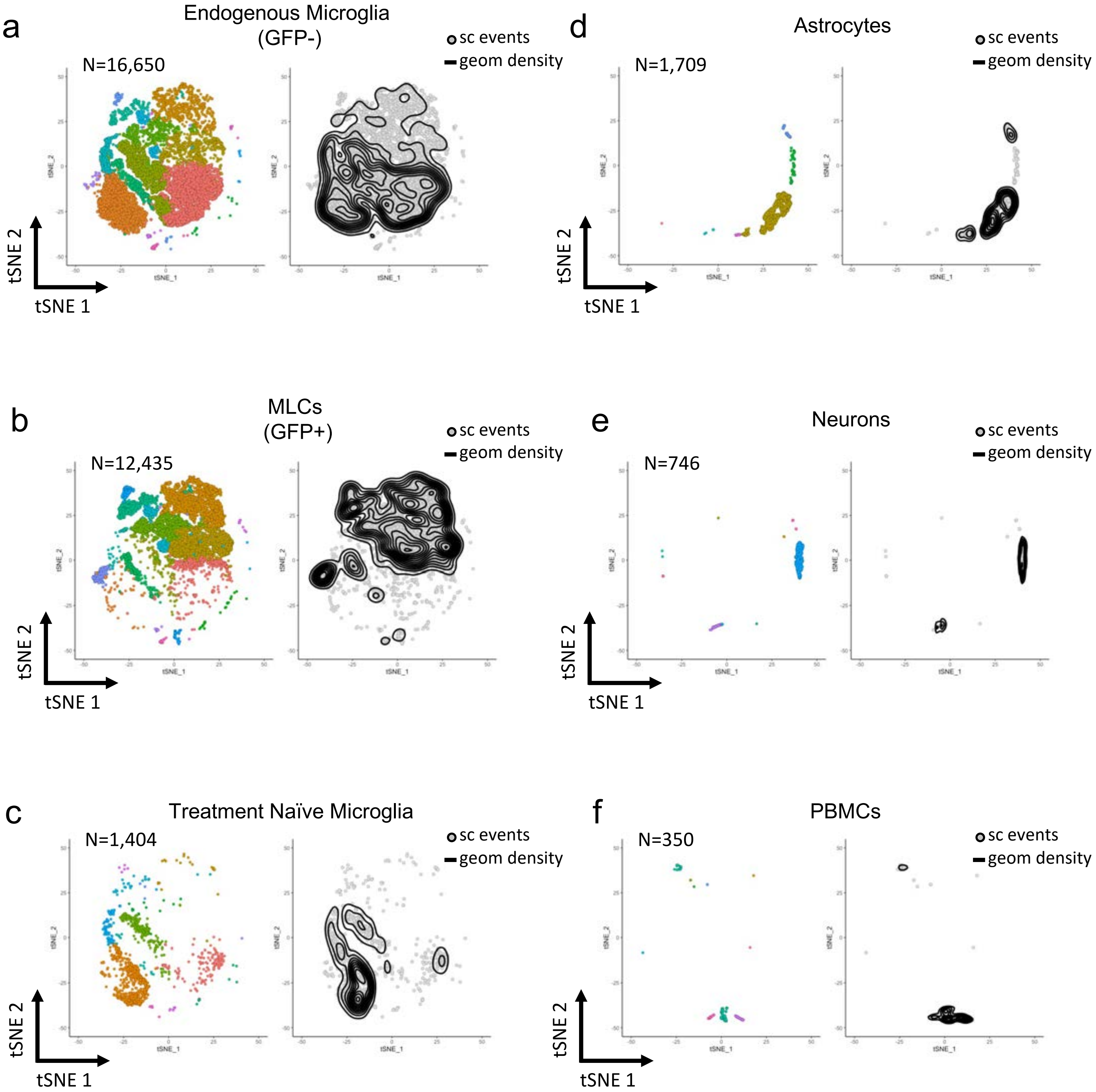


**Supplemental Figure 5. tSNE plots for combined analysis from Figure 3 separated by sample**
Shown are tSNE plots from the analysis shown in figure 3 separated by sample type: endogenous microglia (**a**), MLCs (**b**), treatment naïve microglia (**c**), astrocytes (**d**), neurons (**e**) and PBMCs (**f**).

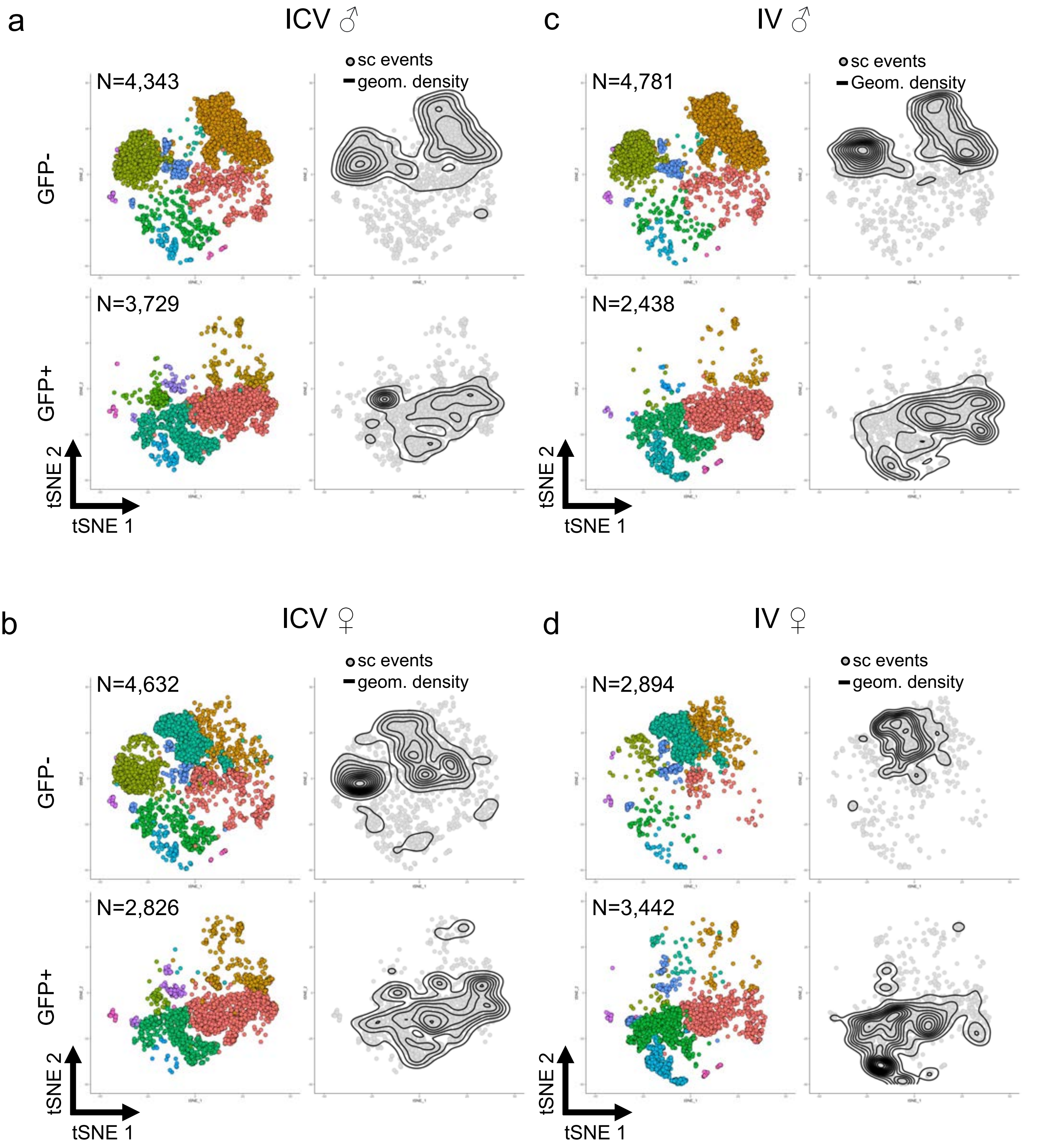


**Supplemental Figure 6. tSNE plots for combined analysis from Figure 4 separated by sample type** Shown are tSNE plots from the analysis shown in figure 4 separated by animal. The GFP+ MLC and GFP- microglia samples are shown for the male (**a**) and female (**b**) dosed via ICV and for the male (**c**) and female (**d**) dosed via IV.

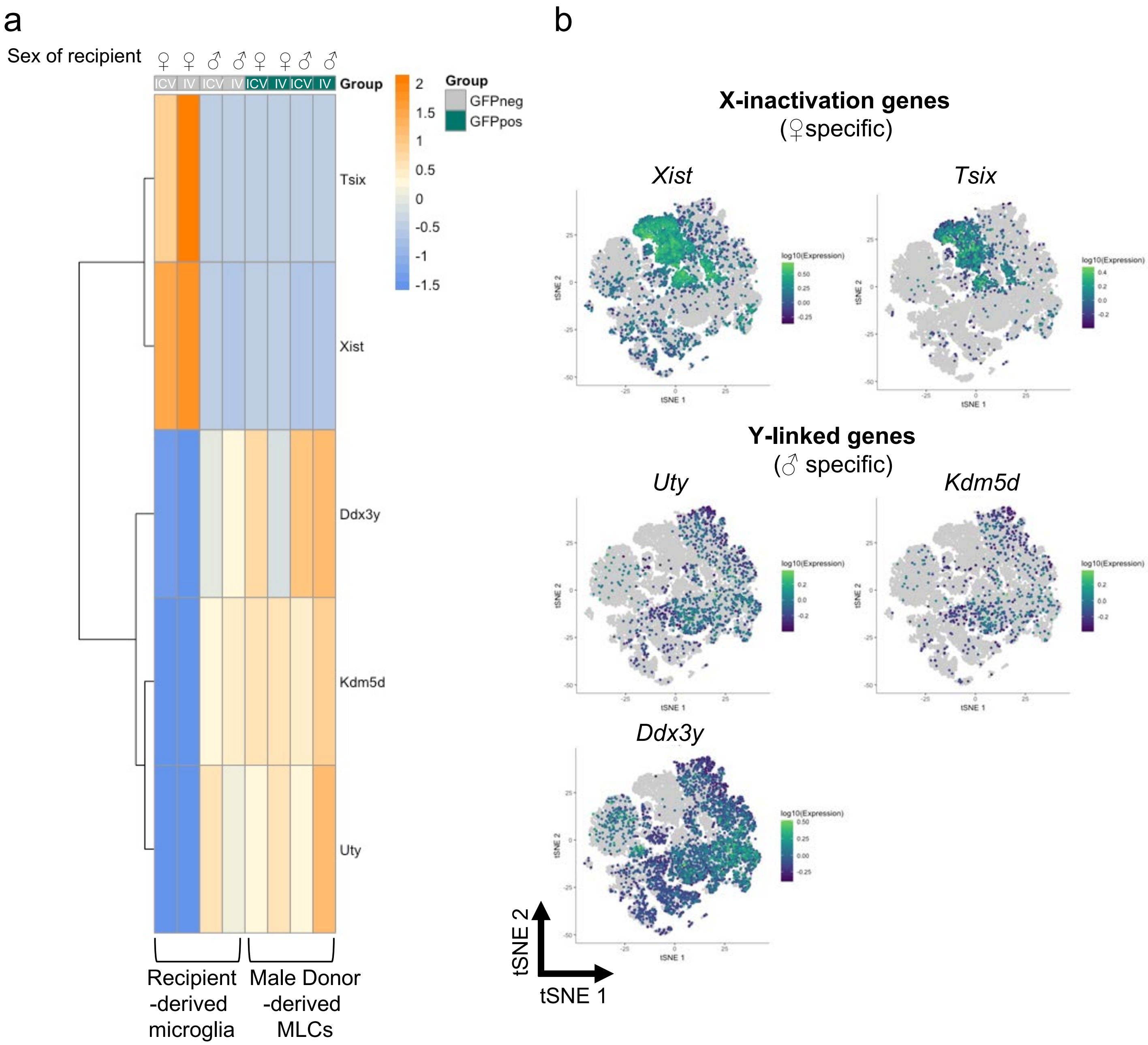


**Supplemental Figure 7. Expression of sex-chromosome genes in MLCs and microglia**
**a** Heat maps of normalized expression for two x-linked genes associated with X-inactivation (*Xist*, *Tsix)* and three Y-linked genes (*Ddx3y*, *Uty*, *Kdm5d)* separated by sample type. **b** tSNE plot colored for expression of the X-linked and Y-linked genes.

a

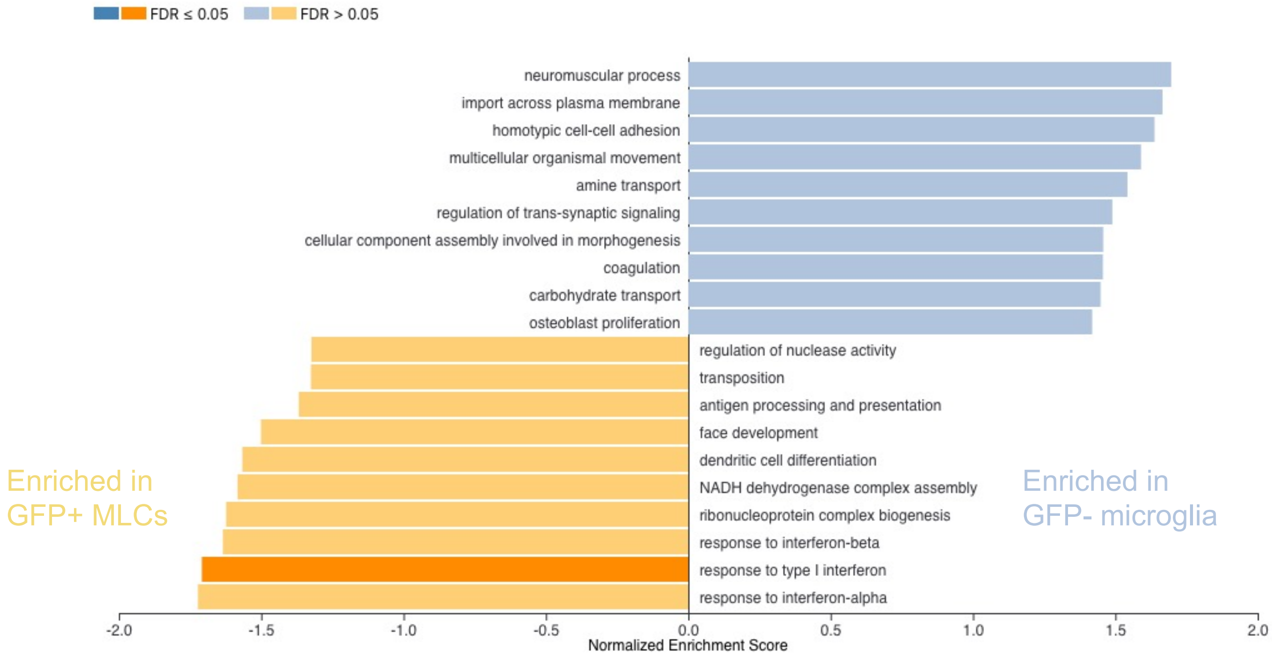


b

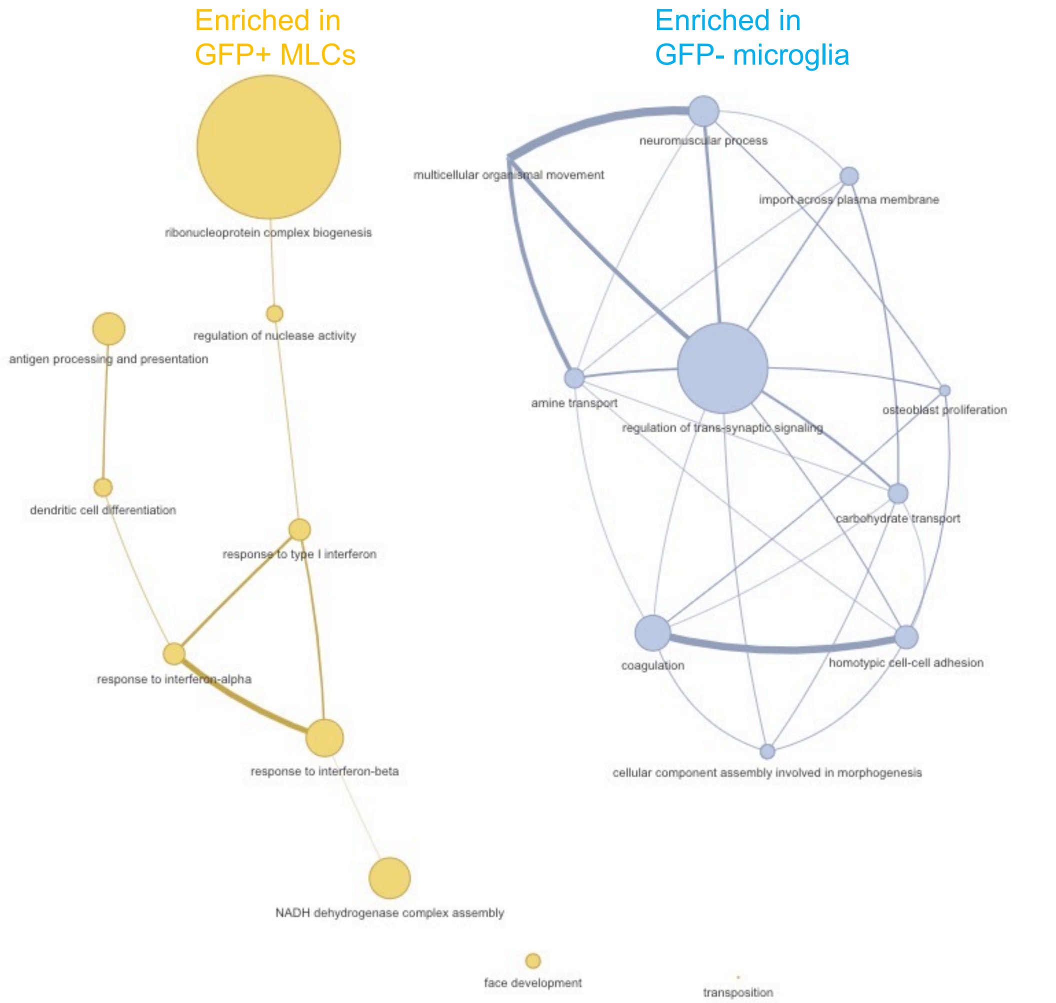


**Supplemental Figure 8. Gene Ontology analysis comparing MLCs to microglia. a** Top ten gene ontology terms associated with genes enriched in MLCS vs. endogenous microglia. GSEAPreranked analysis via webgestalt was used to generate GO term list. **b** GO term pathway analysis visualized for the terms via AutoAnnote (Cytoscape).

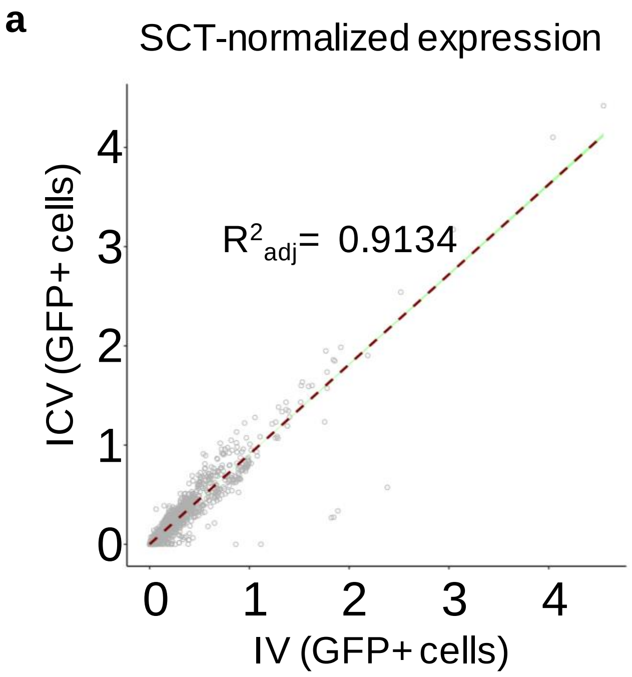


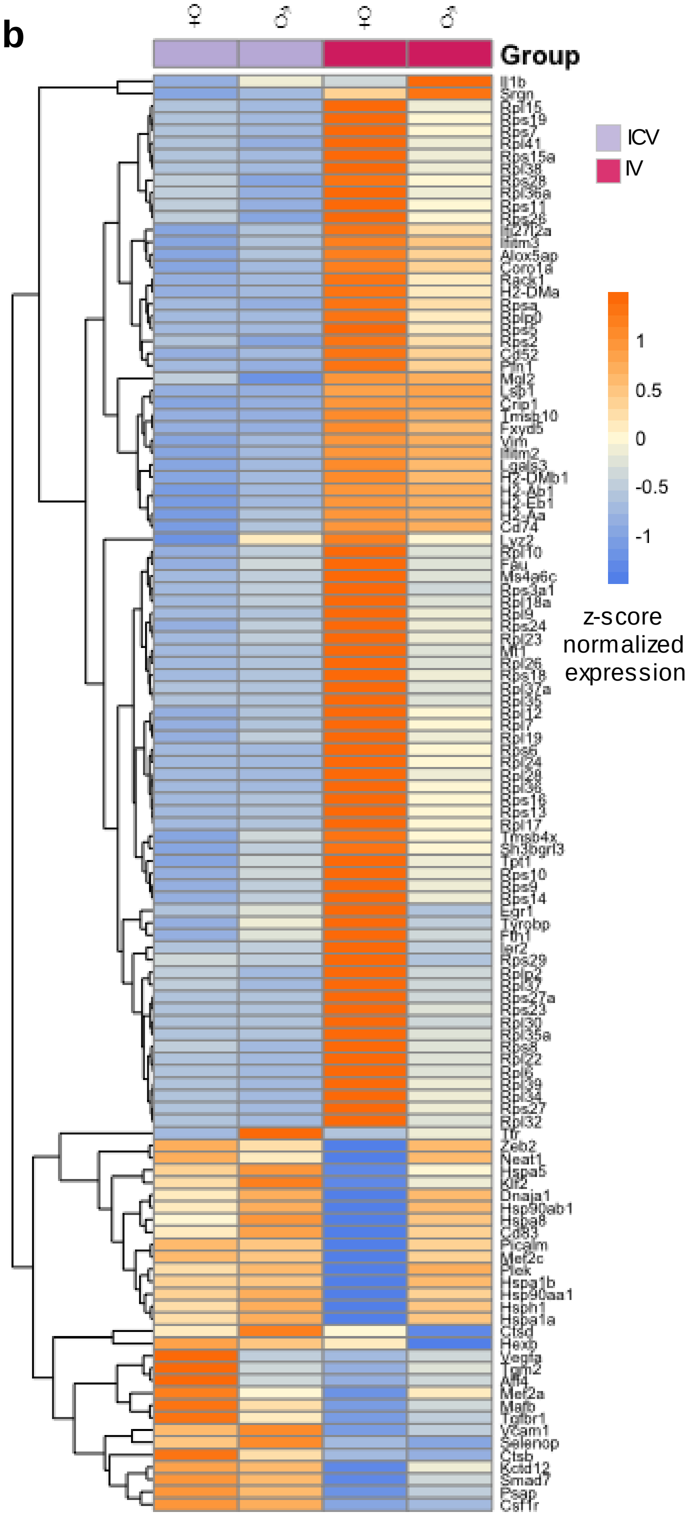


**Supplementary Figure 9. Comparison of IV and ICV-derived MLCs. a** Correlation analysis of MLCs isolated from IV and ICV dosed animals based on normalized single cell gene expression data. **b** Unsupervised clustering based on the global expression profile of each sample type (top). Heatmap showing top differentially expressed genes for each sample type.

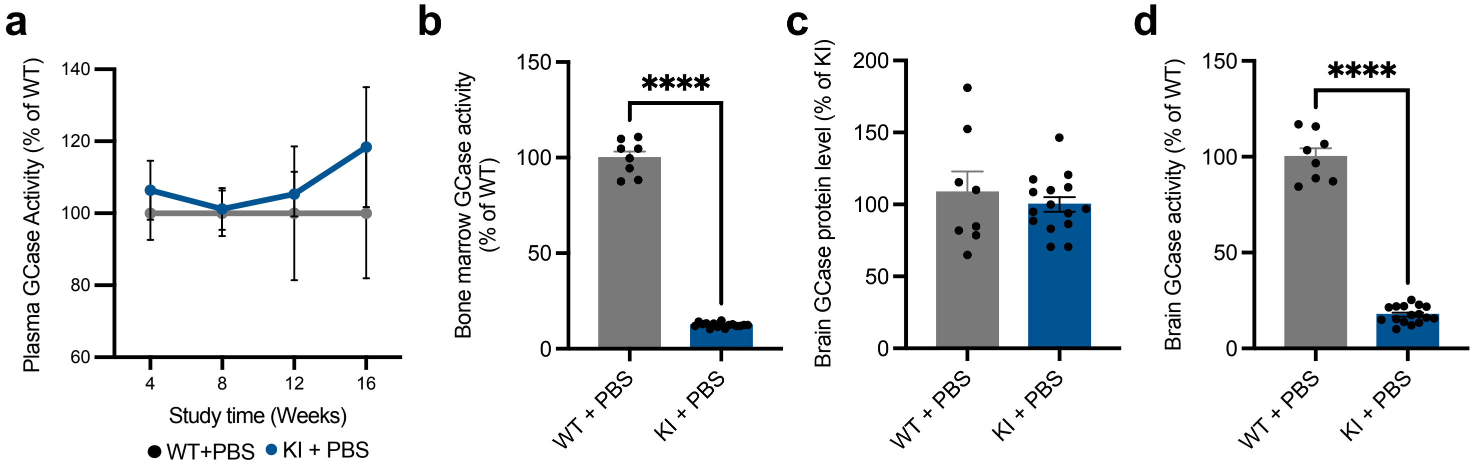


**Supplementary Figure 10. Baseline characterization of homozygous Gba D409V knock-in mouse model.** **a** GCase activity was measured from plasma at four week intervals through the course of the study and normalized to wild-type levels. Lack of difference between wild-type and knock-in animals is likely due to low endogenous activity and background noise. **b** GCase activity (normalized to wild-type animals) at the terminal timepoint of 16-weeks in the bone marrow. **c-d** GCase protein levels (normalized to knock-in animals) and activity (normalized to wild-type animals) were measured in the brain at the terminal timepoint of 16-weeks. T-test was used for statistical analysis. **, $p<0.01$. Bars represent means ± SEM.

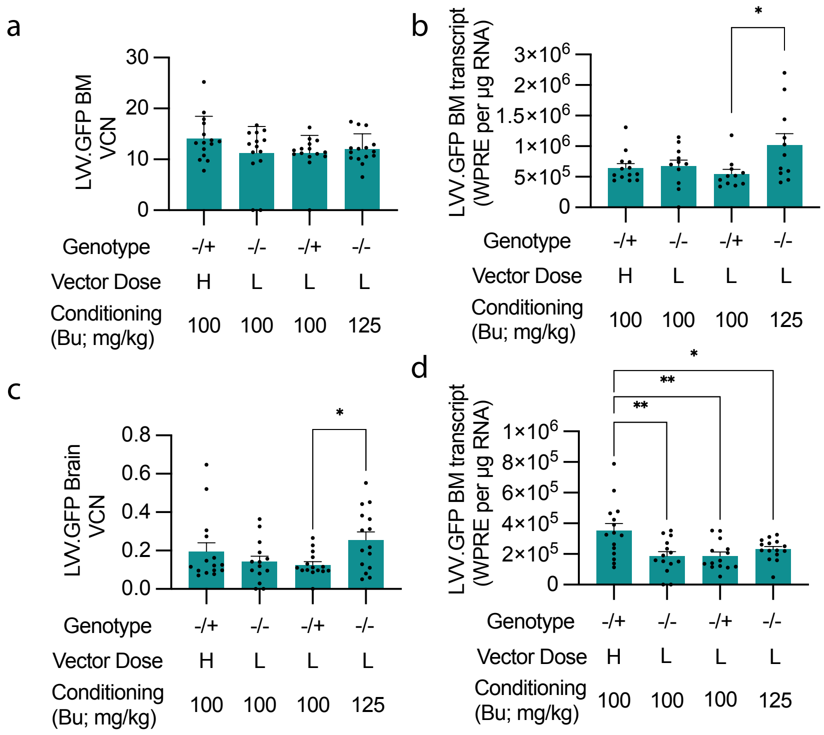


**Supplementary Figure 11. VCN and WPRE quantification for LVV.GFP-treated animals at 16 weeks post-transplantation a** Bone marrow VCN **b** Bone marrow WPRE **c** Brain VCN **d** Brain WPRE**.** All error bars are SEM. Tukey's multiple post-hoc comparison test was conducted for all statistical measures shown, with only significant differences indicated.
